# Why rescue-based transcript ranking can mislead: normalization displacement and shared-control coupling in perturbation transcriptomics

**DOI:** 10.64898/2026.09.27.754762

**Authors:** Hasan M. Jamil, Alexander Gow

**Affiliations:** Department of Computer Science, University of Idaho, 875 Perimeter Drive, Moscow, 83844, ID, United States; Center for Molecular Medicine and Genetics, Wayne State University, 540 E. Canfield Avenue, Detroit, 48201, MI, United States

**Keywords:** perturbation transcriptomics, rescue analysis, statistical calibration, PLP1, PERK, integrated stress response

## Abstract

When a disease-associated transcript moves toward control after a phenotypically effective rescue, the pattern can be read as mediator evidence. We show why this inference can fail even when the computation is reproducible. Across normalization schemes in PLP1-mutant Jimpy brain (GSE277705), the original negative-interaction criterion classified 47.6–76.0% of all expressed genes and 77.6–86.6% of disease-eligible genes. Under total CPM, the excess was driven largely by a +1.016-SD median WT treatment response; median-of-ratios (MOR) reduced it to +0.154 SD while leaving the Jimpy median near zero. Under MOR, mirror enrichment among disease-increased (77.6%) and disease-decreased (74.0%) genes showed selection dependence but could not distinguish mechanical coupling from genuine state restoration. Slope decomposition gave a 48.7% mechanical-to-observed ratio (−0.125 of −0.256); deleting one of four untreated-Jimpy samples moved it from 37.8% to 90.1%. Candidate-local point estimates were 60.8% near TRIB3 and 38.6% near DDIT4; the TRIB3-minus-DDIT4 difference stayed positive in all four deletions (8.6–27.7 points), but its magnitude remained unstable. Oligodendrocyte-specific *Perk* deletion robustly suppressed TRIB3 (*R*^*JP*^ = −0.817, 95% CI [−1.048, −0.587]) and DDIT4 (−0.292, [−0.419, −0.169]), whereas 2BAct resolved neither transcript-specific rescue. Rescue transcriptomics can therefore establish broad state restoration without identifying which downstream transcript mediates it.

## 1. Introduction

A phenotypic rescue accompanied by movement of a disease-associated transcript toward control appears to offer convergent mechanistic evidence: the intervention improves disease and the molecular abnormality moves in the expected direction. That intuition makes rescue-based transcript ranking attractive for candidate prioritization, but it can overstate what the design identifies. A genotype-by-rescue interaction asks whether treatment responses differ by genotype; it does not by itself establish that the disease group moved toward control. In addition, disease induction and apparent disease-group rescue can share the untreated-disease group mean with opposite signs, so selecting large disease effects can mechanically enrich apparently favorable rescue effects. The practical consequence is not merely a biased coefficient. A biologically plausible transcript can rise in a rescue ranking for reasons that are partly generated by the contrast geometry itself.

The statistical ingredients of this problem are not new in isolation. Interaction terms estimate differences between effects and should not be substituted for the corresponding simple effects; errors in interpreting interaction evidence are well documented in neuroscience and related experimental literatures Nieuwenhuis et al. (2011). Mathematical coupling between an initial value and a change score has been recognized since Oldham and repeatedly revisited in biostatistics Oldham (1962); Tu and Gilthorpe (2007). RNA-seq normalization methods such as TMM and median-of-ratios were developed specifically because library-size scaling can fail when composition changes asymmetrically across samples Robinson and Oshlack (2010); Anders and Huber (2010). Cross-fitting is likewise an established data-splitting strategy rather than a new contribution here Chernozhukov et al. (2018). Our contribution is the rescue-specific geometry and an operational adjudication workflow: we derive the shared-control term for disease-versus-rescue contrasts, estimate its abundance-stratified magnitude and sample-level sensitivity in a real factorial rescue design, and pair it with two diagnostics that can be run before interpreting any nominated transcript—a transcriptome-wide threshold-prevalence audit and an arm-wide audit of *R*^*JP*^, *R*^*W T*^, and *I*.

We develop this rescue-aware adjudication framework around two inferential hazards (Fig. 1) and a separate malization requirement. The first safeguard is semantic: report the actual disease-group change under rescue, ^*J P*^, h uncertainty, rather than using the interaction term as a synonym for normalization. The second is statistical: when disease-effect magnitude is used to define a matched background, literal reuse of the same untreated-disease samples should be broken, but any decoupling strategy must itself be audited for cross-cohort transportability or split instability. The normalization safeguard is arm-wide: distinguish denominator-arithmetic controls from informative alternative size-factor models and inspect the distributions of *R*^*JP*^, *R*^*W T*^, and *I*, not only threshold exceedance. The threshold-prevalence audit asks a deliberately simple question before candidate interpretation: if a proposed mechanistic rule classifies a large fraction of the expressed transcriptome, does the rule have enough specificity to rank a nominated mediator at all?

**Figure 1.**
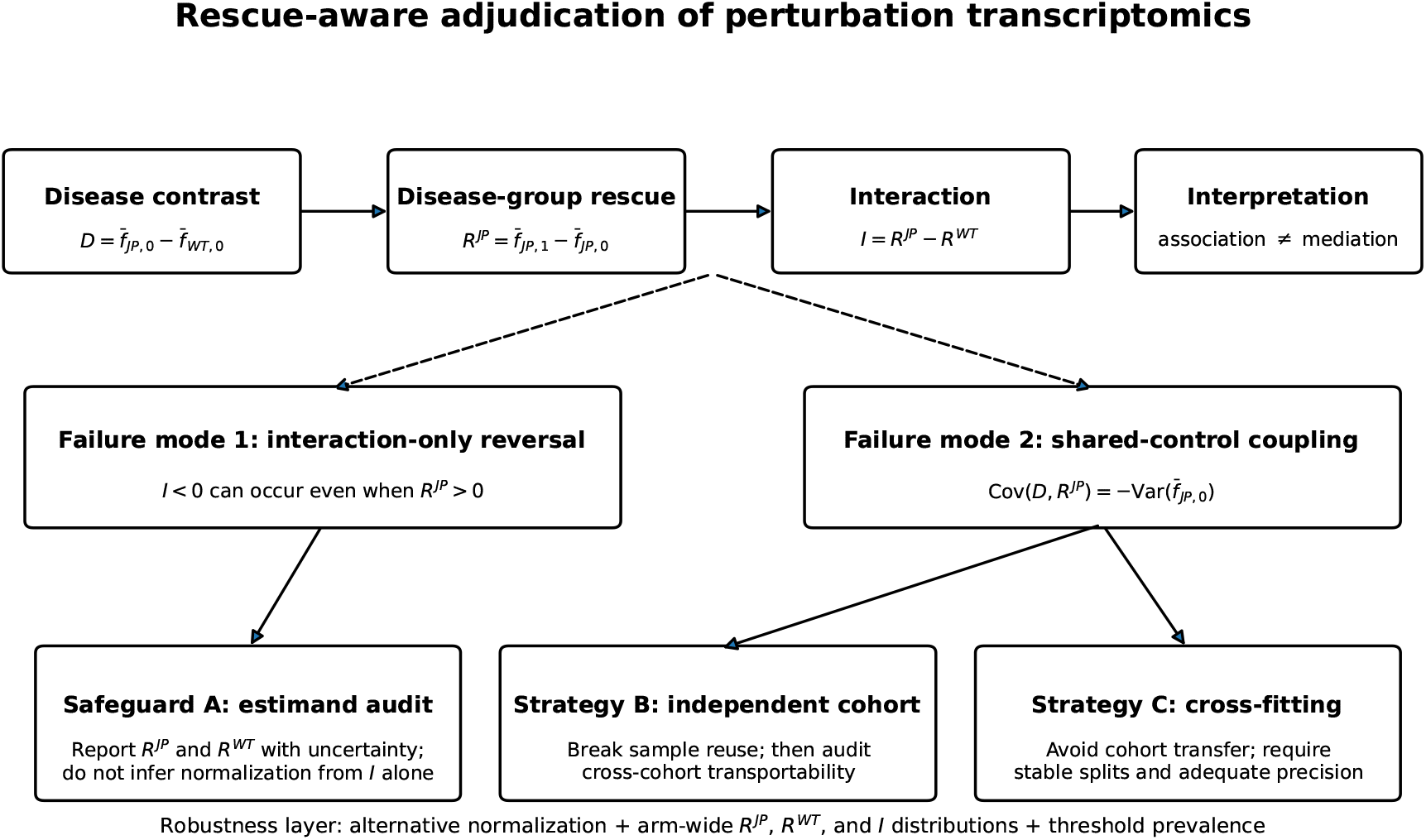
Rescue-aware adjudication design. The framework separates disease induction *D*, the actual disease-group rescue effect *R*^*JP*^, and the genotype-by-rescue interaction *I*. A negative interaction can coexist with an increase in rescued disease animals, while *D* and *R*^*JP*^ are mechanically negatively coupled when they reuse the same untreated-disease mean. Independent-cohort calibration and cross-fitting are conditional decoupling strategies rather than universal corrections: the former requires a transportability audit and the latter sufficient untreated-disease replication for stable partitions. Normalization, arm-wide response distributions, and threshold prevalence form a separate robustness layer. The schematic is methodological and does not encode the empirical outcome of the PLP1 case study.

PLP1-mutant oligodendrocyte disease provides a useful worked case because the upstream rescue node is unusually well constrained. Coding-region *PLP1* mutations can cause proteolipid protein retention in the endoplasmic reticulum (ER), unfolded-protein-response activation, oligodendrocyte loss, and hypomyelination Gow et al. (1998); Southwood et al. (2002); Garbern (2007). Disease severity is allele- and context-dependent Schneider et al. (1992), and recent work has added ER-calcium depletion and impaired COPII trafficking to the pathogenic repertoire Li et al. (2026). The question is therefore not whether mutant PLP1 can create ER stress, but how to distinguish downstream responses that are causal, adaptive, or merely correlated with stress severity.

The PERK/eIF2α integrated stress response (ISR) is central to this question. PERK signaling reduces global translation while enabling selective stress-responsive translation and transcription Pakos-Zebrucka et al. (2016); Han et al. (2013). Yet canonical “pro-death” labels perform poorly in myelinating glia: deleting ATF3 or caspase-12 produces little rescue in PLP1-mutant models, while CHOP can be protective in another PLP1 context Sharma et al. (2007); Sharma and Gow (2007); Southwood et al. (2002); Gow and Wrabetz (2009). Stress induction therefore cannot be equated with death execution.

The Jimpy model adds causal leverage. Elitt and colleagues identified a pathological transition during oligodendrocyte differentiation and showed that Ro 25-6981 can improve oligodendrocyte survival without restoring later myelination Elitt et al. (2018). More recently, Chen et al. (2026) showed that oligodendrocyte-lineage-specific *Perk* deletion reduces caspase-3-positive oligodendrocytes, increases oligodendrocyte survival and myelination, and extends lifespan. Pharmacologic eIF2B activation with 2BAct or ISRIB also improves survival and myelination. These perturbations establish the detrimental role of excessive PERK/ISR signaling in Jimpy disease, but they do not by themselves identify a downstream molecular executor.

Two candidate branches are especially relevant. ATF4 can induce *DDIT4* (REDD1), which can suppress mTORC1 through TSC1/TSC2 Whitney et al. (2009); Kimball and Jefferson (2012); Brugarolas et al. (2004); REDD1 can also support survival in other stress contexts Dennis et al. (2013). ATF4/CHOP can induce *TRIB3*, which has been linked to ER-stress-associated cell death and inhibition of AKT survival signaling Ohoka et al. (2005). Chen et al. (2026) already reported that *Trib3* is induced in Jimpy brain and decreases after oligodendrocyte-specific PERK deletion, nominating a TRIB3/AKT mechanism. A p53-centered mechanism was retained as an exploratory competitor because p53 effects in oligodendrocytes are injury-context dependent Luo et al. (2021); however, the revised analysis treats individual p53-related genes rather than an assumed coherent module.

The motivating counterfactual in this study is documented rather than hypothetical. The first completed classifier used disease induction, a negative interaction threshold, and a minimum reversal fraction and yielded biologically plausible “rescue-consistent” calls. Crucially, under P19 2BAct, DDIT4 could satisfy the interaction-based reversal rule even though its expression increased in treated Jimpy animals. Subsequent versioned audits then exposed shared-control coupling, normalization-dependent arm-wide displacement, abundance dependence, and transportability limitations in independent matching. Because the superseded analyses are retained in the reproducibility archive, the case records how a plausible candidate-level conclusion was weakened not by changing the biology, but by asking whether the rule itself was specific.

If these audits are correct, the biological interpretation of the PMD rescue experiments changes in three ways. First, the P19 2BAct transcriptome does not provide independent transcript-specific corroboration for TRIB3/AKT; the strongest case for TRIB3 rests on disease induction, genetic PERK dependence, prior pathway biology, and a still-needed direct perturbation. Second, DDIT4 warrants direct testing as a robust PERK-responsive comparator, but not as a proven disease mediator because its disease elevation is weak and cohort/normalization dependent. Third, when rescue changes a dominant oligodendrocyte/myelin program in whole brain, bulk transcriptomics may be well suited to confirm broad state restoration while remaining structurally limited for ranking downstream executors. The decisive next experiments are therefore cell-resolved or targeted perturbation/readout studies, not further ranking of the same bulk rescue transcriptome. The contribution of this study is consequently both methodological and biological: it provides a way to audit rescue-based ranking and places an explicit bound on what this experimental format can establish about mechanism.

## 2. Materials and methods

### 2.1. Rescue-adjudication estimands

For a feature *f*, let 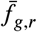 denote mean expression for genotype *g ∈* {*W T, J P* } and rescue status *r ∈* {0, 1}. We define

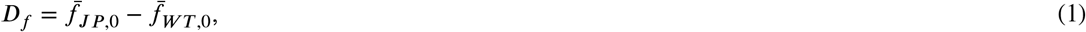

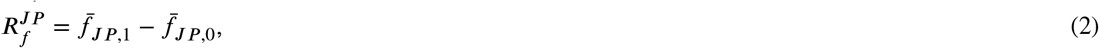

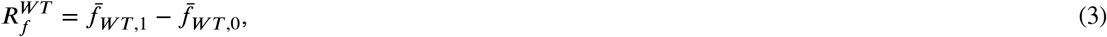

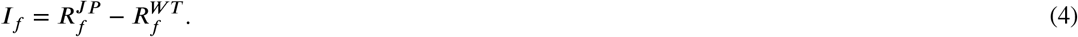

*D*_*f*_ measures disease association, 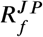 measures what actually happens in rescued disease animals, and *I*_*f*_ measures whether the rescue response differs b^*f*^y genotype.

The original classifier used disease induction, a negative interaction threshold, and a minimum reversal fraction. During the first completed analysis we identified a semantic failure mode: *I*_*f*_ < 0 can occur while 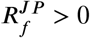 whenever the intervention increases the feature more strongly in WT than in Jimpy. We therefore require the actual disease-group direction to be reported explicitly and do not use a negative interaction as a synonym for normalization.

### 2.2. Shared-control coupling

A second problem arises when genes are selected for large disease induction and then evaluated for rescue using the same untreated-disease samples. Let

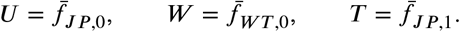

Then *D* = *U − W* and *R*^*JP*^ = *T − U* . If the three group means are independent across biological samples,

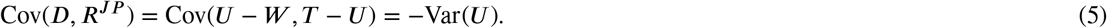

Thus, upward sampling noise in the untreated-Jimpy mean simultaneously increases the estimated disease effect and makes the estimated rescue effect more negative.

We first quantified the practical magnitude of this coupling using the descriptive approximation

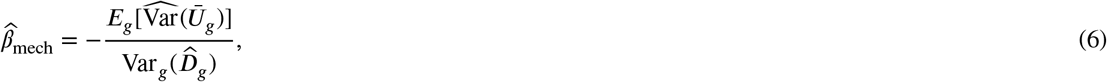

and reported it alongside the observed cross-gene regression slope of *R*^*JP*^ on *D*. Their ratio is not an exact causal decomposition: the approximation treats the average sampling variance of the untreated-disease mean as separable from the across-gene disease-effect distribution. Because RNA-seq variance is expression-dependent, we therefore repeated the calculation within deciles of untreated baseline log-expression and in 500-gene abundance neighborhoods centered on TRIB3 and DDIT4. Candidate-local and expression-stratified values are used when discussing ranking; the transcriptome-wide ratio is retained only as a global design summary.

To quantify sample-level sensitivity without treating genes as independent replicates, we performed a delete-one-sample jackknife over the four untreated-Jimpy P19 2BAct biological samples. Each deletion perturbed only the untreated-disease group that supplies the shared mean *U* ; the WT and treated-Jimpy groups were held fixed. For each deletion we re-estimated non-myelin MOR normalization and recomputed the mechanical slope, observed slope, transcriptome-wide ratio, expression-decile ratios, and the frozen 500-gene candidate-local ratios. We report the full-data estimate and the leave-one-out range as the primary sensitivity summary. Because this design perturbs only the shared-control channel, it isolates sensitivity propagated through *U* rather than bounding total sampling uncertainty in the cross-gene slope. We did not use a naive gene bootstrap as the primary uncertainty analysis because genes within a sample share normalization factors and therefore do not provide independent cross-gene errors.

An initial within-arm matched-*D* analysis is retained in the reproducibility archive for transparency but is not used for inference because it reuses the same untreated-Jimpy samples in both selection and evaluation.

### 2.3. Decoupled specificity calibration

For the primary calibration, disease matching was defined from the independent P19 PERK cohort on the log2CPM scale. The 16 P19 PERK samples and 13 P19 2BAct samples were GSM-disjoint; no accession appeared in both designs. For each candidate, up to 200 non-candidate background genes were selected by proximity in PERK-arm untreated Jimpy-versus-WT disease effect, untreated baseline abundance, and untreated within-genotype residual variability. Candidate rescue behavior was then evaluated only in the separate P19 2BAct cohort. The candidate percentile is the fraction of matched genes whose 2BAct *R*^*JP*^ is less than or equal to the candidate value: 50% is typical, values near 0 are unusually negative, and values near 100 are unusually positive. For descriptive calibration of extreme ranks, we also report (1 + *k*)/(1 + 200), where *k* is the number of matched genes at least as extreme in the relevant tail. This is a finite-sample rank quantity, not a formal *p* value, because matched genes are correlated and two candidates were examined.

Leave-one-out cross-fitting of the four untreated-Jimpy P19 2BAct samples was retained only as a stress test of whether sample splitting could substitute for an independent cohort. Its split-specific percentiles spanned most of the possible range, so cross-fitting at *n* = 4 was deemed non-discriminating and was not used to corroborate candidate-specific inference.

### 2.4. Genome-wide threshold prevalence and normalization audit

To determine whether the original interaction rule had arm-specific discriminating power, we applied the frozen threshold *I ≤ −*0.25 SD to one fixed expressed-gene universe in the P19 2BAct cohort. We report the proportion of all expressed genes meeting the interaction threshold, the proportion among genes satisfying the original disease-induction requirement, the proportion satisfying the original interaction-plus-reversal-fraction rule, and the proportion satisfying the post hoc strict rule that additionally requires *R*^*JP*^ < 0. As a symmetric selection-dependence diagnostic, we also selected strongly disease-decreased genes (*D ≤ −*0.25 SD) and evaluated the mirror criteria *I ≥* +0.25 SD, the same reversal-fraction threshold, and *R*^*JP*^ > 0. The mirror was designed to test whether enrichment reverses with the sign of disease-effect selection; mechanism-specific attribution to shared-control coupling was assessed separately by the slope decomposition.

Because rescue changes large expression programs in Jimpy brain, we also audited normalization. Three schemes were compared: (i) the original total-library CPM; (ii) a denominator-arithmetic control in which the frozen set *Mbp, Plp1, Mag, Mog, Mobp, Klk6, Opalin*, and *Ermn* was excluded only from the CPM library-size denominator; and (iii) a DESeq-style median-of-ratios normalization whose size factors were estimated only from non-myelin genes with positive counts in every sample of the arm. Candidate counts themselves were never excluded. For a myelin-set read fraction *f*, excluding the set from the denominator shifts every retained transcript by − log_2_(1 *− f* ); therefore the expected change in a rescue contrast is determined arithmetically by the difference in *f* between treated and untreated cells. This makes scheme (ii) a leverage check rather than an independent biological sensitivity analysis. Scheme (iii) is the informative alternative normalization because its size factors are estimated from the broader non-myelin count distribution. After the arm-wide sanity audit showed that total CPM produced large positive median WT responses in both P19 arms whereas MOR substantially recentered them, MOR was designated primary for genome-wide interaction-distribution and prevalence inference; total CPM is retained as the frozen original analysis and candidate-level sensitivity comparison. The expressed-gene universe was held fixed from the original total-CPM criterion so that normalization could not change the genes being audited. We summarized the median, interquartile range, and directional fractions of *R*^*JP*^, *R*^*W T*^, and *I* for every P19 arm and normalization, together with MOR size factors, raw library sizes, and myelin-set read fractions by genotype-by-rescue cell. These quantities identify which side of the interaction drives a shift but do not, by themselves, assign a biological cause to the normalization difference.

### 2.5. Datasets and preprocessing

Five public datasets were retained from the original hypothesis-directed analysis. GSE277705 is the load-bearing rescue dataset and contains 57 bulk-brain RNA-seq samples from the Jimpy rescue study of Chen et al. (2026). The P19 genetic arm contains four biological replicates in each of four cells: WT Cre−, WT Olig2-Cre+, Jimpy Cre−, and Jimpy Olig2-Cre+. The P19 2BAct arm contains WT control *n* = 3, WT 2BAct *n* = 3, Jimpy control *n* = 4, and Jimpy 2BAct *n* = 3. GSM-level manifest auditing confirmed that the 16 P19 PERK samples (GSM8527867–GSM8527882) and 13 P19 2BAct samples (GSM8527911–GSM8527923) are disjoint, with zero shared accessions. P7 and P14 2BAct arms were retained for descriptive temporal context.

Raw GSE277705 counts were mapped using a pinned GRCm38/Ensembl release-91 annotation. Each intervention arm was normalized independently to counts per million and transformed as log_2_(CPM + 0.5). The original analysis also standardized each gene within arm; those z-scale results are retained as sensitivity analyses, but log2CPM is primary for paired between-gene contrasts because subtracting effects expressed in different gene-specific SD units has no common physical scale.

GSE111595 contains wild-type and *Plp* ^jimpy^ OPCs and day-1 oligodendrocytes with three biological replicates per genotype-by-state cell Elitt et al. (2018); Elitt and Tesar (2018). GSE81792 contains rat OPCs treated with 0 or 5 *µ*M tunicamycin for 2 or 8 h, with three replicates per treatment-by-time cell; both time points precede overt cell death in the source experiment Hetman et al. (2016). These datasets provide temporal and pre-death context only. Two human single-nucleus datasets, GSE180759 and GSE118257, were retained as external context for stress-program heterogeneity Absinta et al. (2021a,b); Jäkel et al. (2019a,b); they are not used as causal evidence for PMD.

### 2.6. Candidate definitions and program coherence

TRIB3 and DDIT4 were the primary individual-gene candidates. A p53-related set (*Cdkn1a, Mdm2, Pmaip1, Gadd45a, Bbc3, Bax*) was retained for exploratory gene-level analysis. Previously defined multi-gene stress programs were demoted to descriptive summaries after a coherence audit showed that several did not behave as stable latent variables across rescue arms. Coherence diagnostics included median pairwise Spearman correlation, fraction of positive pairwise correlations, PC1 variance explained, and constituent-gene sign concordance. The myelin/differentiation program served as a positive biological control because it was highly coherent in both P19 arms.

RNA abundance was not treated as a direct measurement of AKT or mTORC1 biochemical activity. Likewise, CASP3 and related apoptosis transcripts were treated as transcriptional readouts rather than measurements of caspase activation or apoptosis execution.

### 2.7. Uncertainty and exact tests

For prespecified candidate genes, biological replicates were resampled within the four genotype-by-rescue cells for 20,000 nonparametric bootstrap replicates, yielding percentile 95% intervals for *D, RJP*, and *I*. Exact rescue-label permutation tests were computed within genotype. At P19, the PERK Jimpy comparison has 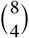 = 70 possible assignments and the 2BAct Jimpy comparison has 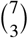 = 35.

Because TRIB3 and DDIT4 are measured in the^3^same animals, their primary paired contrast was

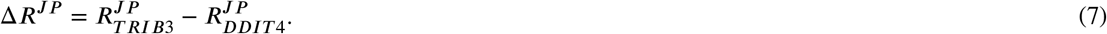

The paired bootstrap resampled the same sample identifiers for both genes. The log2CPM contrast is primary; the z-scale contrast is reported only as sensitivity.

Thirty individual frozen genes entered the gene-level Benjamini–Hochberg correction in each P19 arm. Exact candidate tests are reported separately because the principal candidate comparisons were prespecified.

### 2.8. Cross-arm precision and composition sensitivity

The two P19 designs are separate, GSM-disjoint cohorts and were never pooled for normalization. To quantify empirical cross-arm variability, we compared untreated Jimpy-versus-WT log2CPM effects for the 30 frozen genes and summarized the distribution of absolute between-arm differences. This is an empirical reproducibility audit, not a formal measurement-error floor. Because the normalization audit showed that DDIT4’s 2BAct-arm disease effect is normalization-sensitive, cross-arm magnitude comparisons are interpreted only together with the alternative-normalization results.

GSE277705 is bulk whole-brain RNA-seq. As a diagnostic sensitivity analysis, a myelin/oligodendrocyte expression proxy was formed from *Mbp, Mag, Mog, Mobp, Klk6, Opalin*, and *Ermn*; *Plp1* was excluded. Candidate models were refit as *y ∼ J P* + *rescue* + *J P* × *rescue* + *myelin*_*proxy*. Because the proxy is treatment-responsive and may lie on the causal pathway, adjusted estimates are not interpreted causally.

### 2.9. Reproducibility policy

The analysis was versioned rather than silently overwritten. The first completed classifier, the post hoc strict-direction correction, the v0.2 matched-*D* analysis, the diagnosis of its shared-control coupling problem, the decoupled v0.3 analyses, the v0.4 normalization audit, the v0.5 abundance-stratified coupling audit, the v0.6 interaction-distribution/size-factor diagnostic, and the v0.7 symmetric prevalence diagnostic are all retained in Supplementary Data S1. Analyses added after external review were frozen before rerunning expression and are labeled as robustness analyses rather than preregistered tests. Version 0.9 additionally narrowed the interpretation of the mirror diagnostic, distinguished selection-conditioned symmetry from the slope-based coupling estimate, and made explicit the structural insulation of *R*^*JP*^ from WT-specific normalization displacement. Version 0.10 adds only the prespecified sample-deletion sensitivity analysis for the coupling estimator and corresponding editorial corrections.

## 3. Results

The case study was analyzed as an audit of a frozen rule rather than as a fresh candidate search. We therefore ask first whether the rule discriminates against the expressed-transcript background, then diagnose the arm-wide and shared-control mechanisms that determine its prevalence, and only then return to TRIB3, DDIT4, and exploratory candidates. This order separates evidence that the intervention changes disease from evidence that any particular transcript uniquely tracks that rescue.

### 3.1. The frozen rescue criterion classifies much of the transcriptome

The original total-CPM analysis and the MOR analysis expose different components of the same prevalence problem. Among the fixed 17,684 expressed genes, 75.6% met *I ≤ −*0.25 SD under total CPM, 76.0% under the denominator-only non-myelin CPM control, and 47.6% under non-myelin MOR. Among disease-eligible genes (*D ≥* 0.25 SD), the corresponding fractions were 85.9%, 86.6%, and 77.6%. The original interaction-plus-reversal-fraction rule classified 85.4%, 86.1%, and 77.1% of disease-eligible genes, while the strict rule that additionally required *R*^*JP*^ < 0 classified 54.3%, 54.7%, and 58.8%, respectively (Fig. 2). MOR is primary for arm-wide prevalence interpretation because it substantially recenters the treatment-wide WT displacement documented below; total CPM is retained as the frozen original analysis.

**Figure 2.**
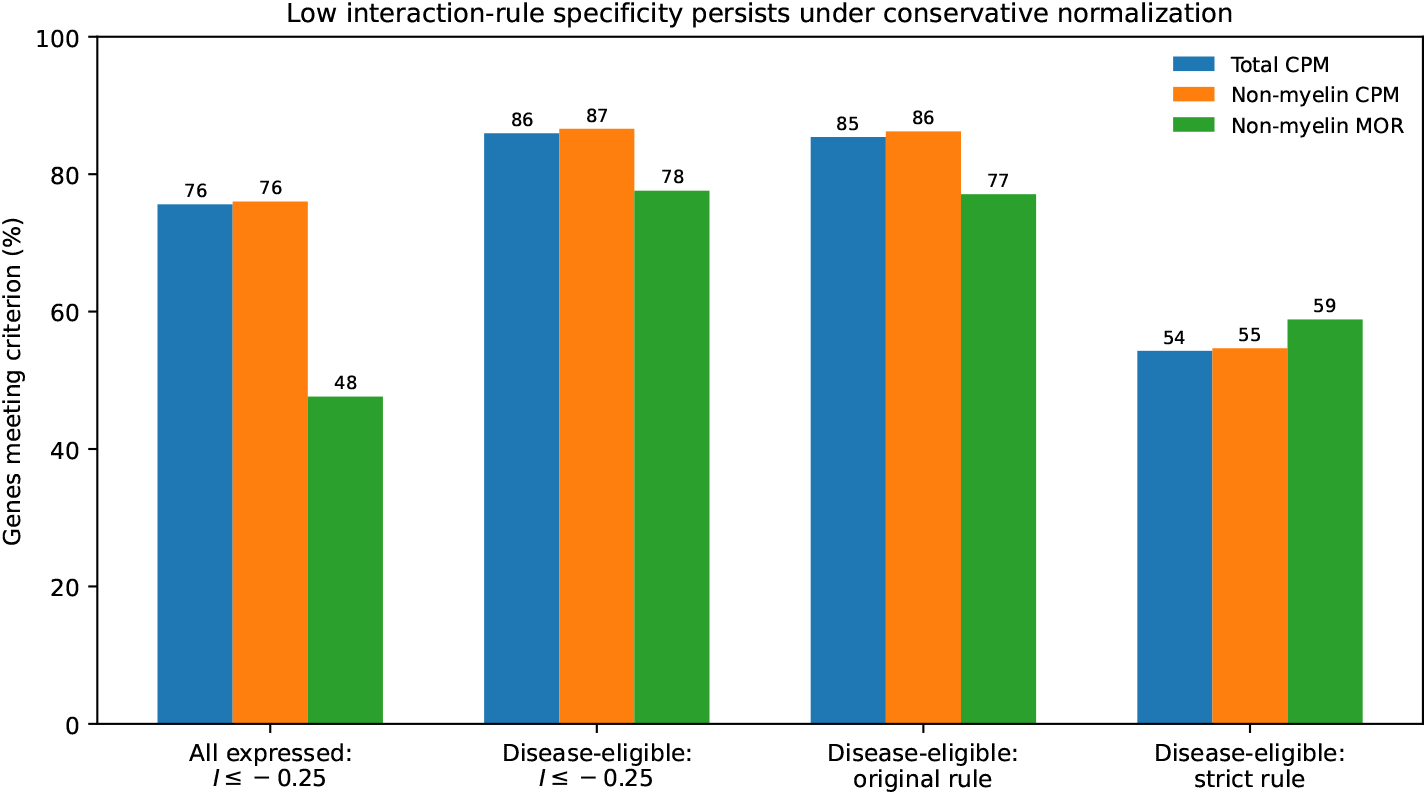
Genome-wide prevalence of the frozen P19 2BAct rescue criteria under three normalization schemes. MOR is primary for arm-wide interaction inference; total CPM is retained as the frozen original analysis and non-myelin CPM is a denominator-arithmetic control. Under MOR, 47.6% of all expressed genes but 77.6% of disease-eligible genes still meet the negative-interaction threshold, motivating the symmetric selection-dependence diagnostic in the text and Supplementary Table S1E.

The total-CPM excess is transcriptome-wide and is driven primarily by the WT side of the interaction. Under total CPM, the median P19 2BAct Jimpy response was essentially zero (*R*^*JP*^ = −0.002 SD; IQR [−0.261, 0.338]), whereas the median WT response was strongly positive (*R*^*W T*^ = +1.016 SD; IQR [0.493, 1.348]). The resulting interaction distribution had median *I* = −1.014 SD (IQR [−1.476, −0.278]), with 81.2% of genes below zero. Under MOR, the Jimpy median remained near zero (+0.003 SD), but the WT median was attenuated to +0.154 SD and the interaction median to −0.149 SD; 53.4% of interactions were negative. Thus, the extreme all-gene prevalence under CPM reflects a normalization-dependent arm-wide WT displacement rather than broad transcriptomic reversal in Jimpy.

The residual MOR enrichment among disease-selected genes has two nonexclusive explanations: shared-control coupling and genuine broad reversal of the disease state. Of 8,335 genes with *D ≥* 0.25 SD, 77.6% had *I ≤ −*0.25 SD. The prespecified mirror test selected 6,806 genes with *D ≤ −*0.25 SD: 74.0% had *I ≥* +0.25 SD. The interaction-plus-reversal-fraction prevalences were similar rather than identical (77.1% versus 72.8%), whereas the strict direction-consistent rule was nearly symmetric (58.85% versus 58.77%). This mirror pattern does not uniquely identify shared-control coupling, because genuine partial restoration of the Jimpy transcriptional state predicts the same sign reversal. Its narrower diagnostic value is that the enrichment changes direction with disease-effect selection and therefore is not well explained by a one-sided threshold-scaling artifact. Attribution to shared-control coupling instead rests on the cross-gene slope decomposition below, which quantifies the component expected from reuse of the untreated-disease mean. Importantly, the specificity conclusion is mechanism-robust: whether the remaining gradient reflects genuine state restoration or other biology, a rule satisfied by most disease-selected genes cannot by itself rank a nominated mediator (Supplementary Table S1E).

The PERK arm shows the same normalization lesson at the transcriptome level even though its candidate-level rescue estimates are stable. Under total CPM, median *R*^*JP*^ = +0.016 SD, median *R*^*W T*^ = +0.562 SD, and median *I* = −0.525 SD; under MOR the corresponding medians are +0.032, +0.036, and +0.049 SD. This contrast is structurally informative: *R*^*JP*^ depends only on treated and untreated Jimpy samples and therefore cannot inherit a displacement confined to WT samples, whereas *R*^*W T*^ and consequently *I* = *R*^*JP*^ *− R*^*W T*^ can. That insulation explains why the candidate-level *R*^*JP*^ estimates remain stable while interaction-based prevalence is normalization-sensitive. It also explains why the strict-rule prevalence need not move with the interaction prevalence: from total CPM to MOR, disease-eligible *I ≤ −*0.25 falls from 85.9% to 77.6%, while the rule that additionally requires the Jimpy-specific direction *R*^*JP*^ < 0 rises from 54.3% to 58.8%. Full *R*^*JP*^, *R*^*W T*^, and *I* distributions are reported in Supplementary Table S1C. The table reports separate gene-wise marginal medians; because medians are nonlinear, median *I* need not equal median *R*^*JP*^ minus median *R*^*W T*^ even though *I* = *R*^*JP*^ *− R*^*W T*^ holds exactly for every gene.

The frozen eight-gene myelin set was markedly depleted in Jimpy relative to WT but still represented a small percentage of total reads (Table 1). In the PERK arm it increased from 0.0838% to 0.2040% in Jimpy and from 0.7971% to 0.9774% in WT; in the 2BAct arm it increased from 0.0838% to 0.1211% in Jimpy and from 0.9243% to 1.4633% in WT. Direct raw-count recomputation confirmed that the two independently sampled untreated-Jimpy means merely round to the same value: 0.083776% in PERK and 0.083822% in 2BAct. By contrast, the nominally analogous untreated-WT cells differed from 0.7971% to 0.9243%, a 16.0% relative difference, providing an independent illustration of cross-cohort variability. Excluding the set from the CPM denominator is predicted to alter *R*^*JP*^ by only 0.00174 log2 units in PERK and 0.00054 in 2BAct, while the corresponding WT changes are 0.00262 and 0.00787. The near-identity of total and non-myelin CPM is therefore arithmetically forced and carries little evidential weight. In P19 2BAct, median MOR size factors changed from 1.276 to 1.368 in WT and from 1.149 to 1.182 in Jimpy, whereas raw-library-size medians changed by similar but not identical proportions (Supplementary Table S1D). These cell-level differences are consistent with MOR attenuating the large WT-centered CPM shift, but they do not uniquely identify its biological or technical source.

**Table 1.** Frozen eight-gene myelin-set fraction of total reads in all P19 genotype-by-rescue cells. The two GSM-disjoint untreated-Jimpy means round coincidentally to 0.0838%; direct recomputation gives 0.083776% (PERK) and 0.083822% (2BAct).

| Arm | WT control | WT rescue | Jimpy control | Jimpy rescue |
| --- | --- | --- | --- | --- |
| PERK deletion | 0.7971% | 0.9774% | 0.0838% | 0.2040% |
| 2BAct | 0.9243% | 1.4633% | 0.0838% | 0.1211% |

### 3.2. Shared-control coupling is substantial in point estimates but imprecisely resolved across abundance strata

The transcriptome-wide shared-control point estimate was large enough to matter empirically, but its magnitude is precisely resolved at four untreated-Jimpy replicates. In the P19 2BAct arm under total CPM, the observed cross-gene slope of *R*^*JP*^ on *D* was −0.291 and the estimated mechanical contribution from reuse of the untreated-Jimpy mean was −0.142 (48.9% of the observed negative slope magnitude). Under non-myelin MOR the corresponding full-data values were −0.256 and −0.125, giving a 48.7% mechanical-to-observed ratio. Across the four delete-one-sample recomputations, the observed slope ranged from −0.341 to −0.217, the mechanical slope from −0.202 to −0.129, and the descriptive ratio from 37.8% to 90.1%. Although every leave-one-out observed slope remained negative, a conventional normal-theory jackknife sensitivity interval for that unbounded slope spans zero ([−0.589, +0.077]); with only four untreated-Jimpy samples we therefore do not treat the denominator gradient itself as statistically resolved. The 48.7% value should be read as a descriptive ratio, not a bounded fraction with a precise confidence interval. The remaining slope can include genuine disease-state reversal as well as other cross-gene biological or statistical structure. The abundance-stratified audit likewise shows that a single global percentage should not be treated as universal. nder non-myelin MOR, full-data point estimates of the mechanical-to-observed ratio ranged from 26.8% to 67.8% across baseline-expression deciles in the 2BAct arm (Fig. 3), but the series was non-monotonic and sample-deletion sensitivity was wide. In 500-gene abundance neighborhoods centered on the candidates, the full-data ratio was 60.8% around TRIB3 and 38.6% around DDIT4. Their paired difference was 22.2 percentage points, and all four delete-one estimates remained positive, ranging from 8.6 to 27.7 points. The ordering is therefore sign-stable across the four deletions, but the size of the TRIB3–DDIT4 gap is not stable enough at *n* = 4 to support a resolved quantitative separation.

**Figure 3.**
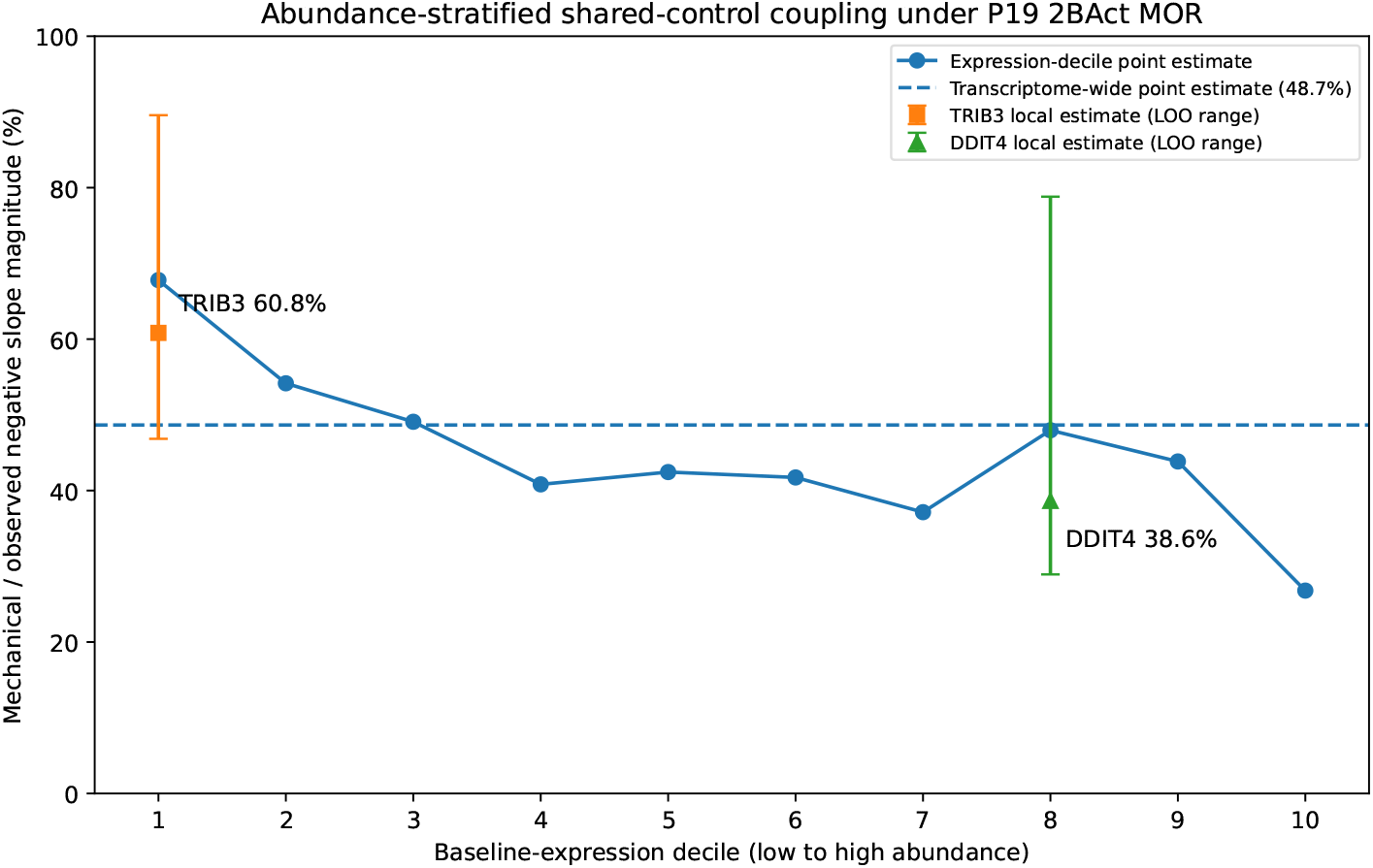
Abundance-stratified point estimates of shared-control coupling in P19 2BAct under non-myelin MOR. The line shows the descriptive mechanical-to-observed negative-slope ratio within baseline-expression deciles; the dashed reference is the transcriptome-wide point estimate (48.7%). Candidate-local points summarize 500-gene abundance neighborhoods around TRIB3 (60.8%) and DDIT4 (38.6%); vertical bars show the range across four delete-one untreated-Jimpy recomputations. The local ranges overlap substantially, although the paired TRIB3-minus-DDIT4 difference remained positive in all four deletions (8.6–27.7 percentage points; Supplementary Table S1F).

These values remain descriptive approximations rather than causal partitions, but they directly quantify the design artifact at the abundance relevant to candidate ranking. TRIB3 itself lies in the lowest baseline-expression decile in the 2BAct arm: under MOR its untreated baseline is approximately 0.25 on the log_2_(CPM + 0.5) scale (about 0.69 CPM), while the decile median is 0.558 (about 0.97 CPM). DDIT4 lies in decile 8. This abundance separation is consistent with TRIB3’s larger local coupling point estimate, greater cross-cohort disease-effect variability, and wide 2BAct bootstrap interval, but the jackknife does not resolve the local coupling difference from DDIT4. The paired log2CPM contrast remains well defined, but it compares transcripts with substantially different precision and dynamic range and is therefore not treated as equal-precision evidence of mechanistic separation.

### 3.3. What survives biologically: PERK deletion suppresses TRIB3 and DDIT4

The strongest within-arm biological result remained the oligodendrocyte-lineage-specific PERK perturbation. Under the original total-CPM normalization, TRIB3 decreased by *R*^*JP*^ = −0.819 log2CPM (95% bootstrap CI [−1.038, −0.599]; exact one-sided *p* = 0.0143), while DDIT4 decreased by −0.295 ([−0.395, −0.187]; *p* = 0.0143). The informative non-myelin median-of-ratios (MOR) sensitivity produced essentially the same rescue effects: TRIB3 *R*^*JP*^ = −0.817 ([−1.048, −0.587]) and DDIT4 −0.292 ([−0.419, −0.169]), with the same exact *p* values (Fig. 4;Table 2).

**Table 2.**
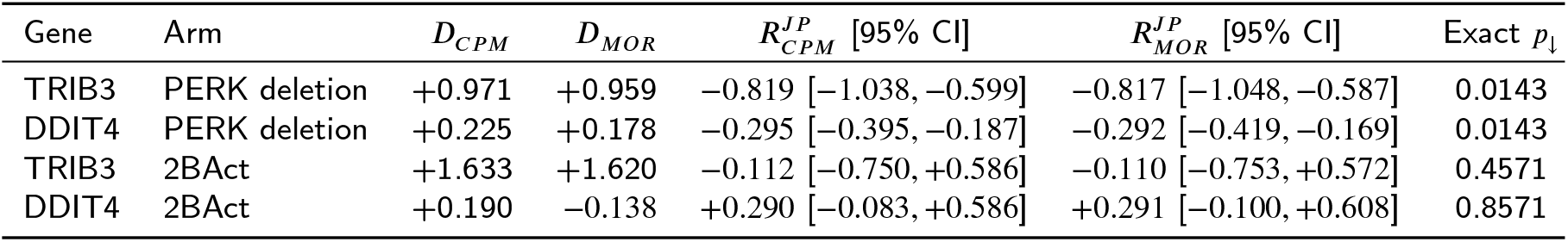
Primary P19 candidate estimates. MOR denotes non-myelin median-of-ratios normalization. The non-myelin CPM denominator control is omitted because its leverage is arithmetically constrained by the small frozen myelin-set read fraction.

**Figure 4.**
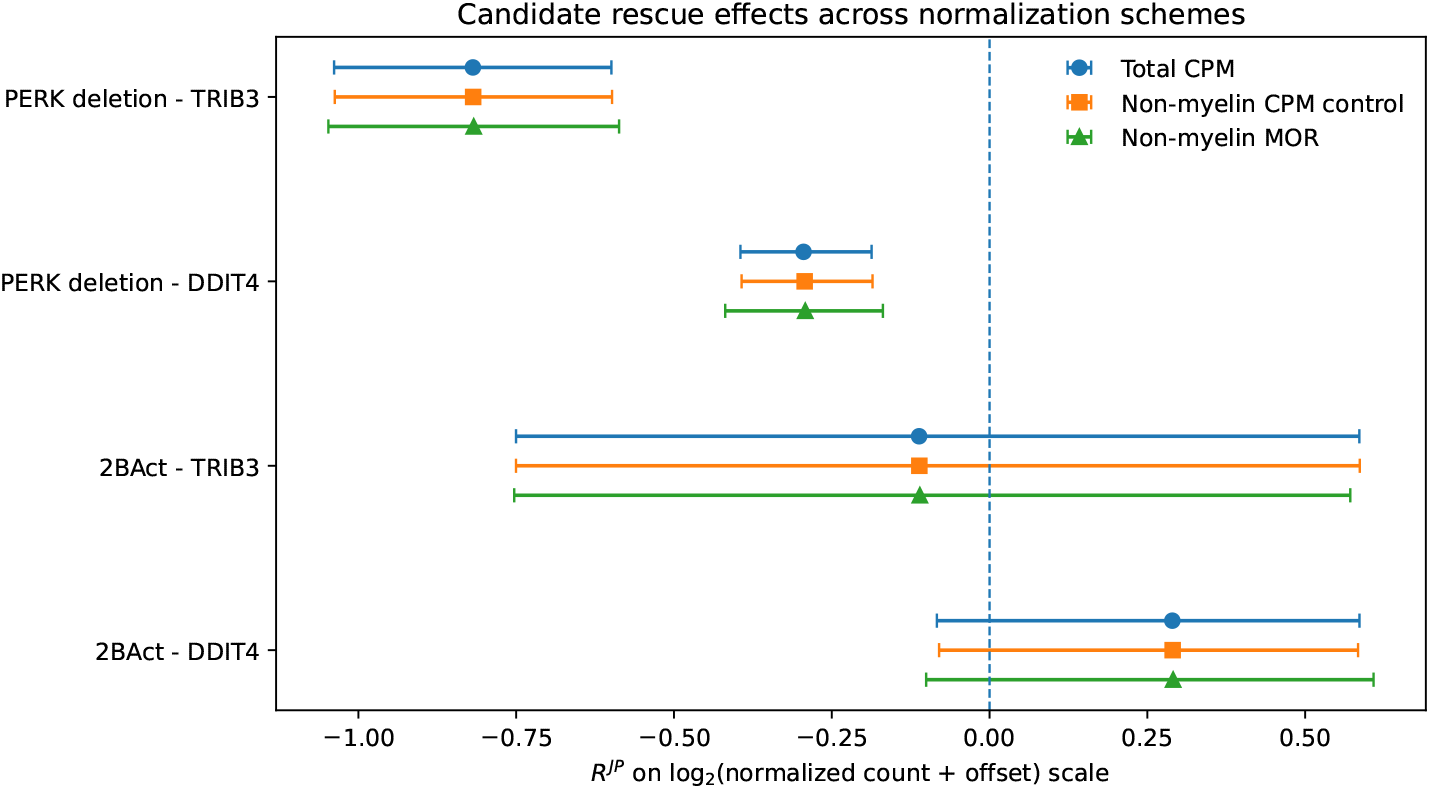
TRIB3 and DDIT4 disease-group rescue effects under three normalization schemes. Each normalization has one consistent legend entry across all four candidate-by-arm rows. Points show *R*^*JP*^ and bars show 20,000-resample bootstrap 95% intervals. Total CPM and non-myelin MOR provide the principal comparison; non-myelin CPM is an arithmetic denominator control with intrinsically small leverage for the frozen eight-gene set. Both genes decrease robustly after oligodendrocyte-specific PERK deletion; neither P19 2BAct effect is resolved from zero.

The denominator-only non-myelin CPM control changed the PERK Jimpy rescue contrast by only the amount ated by its small change in frozen myelin-set read fraction (expected denominator contribution 0.00174 log2 units). The resulting candidate shifts of less than 0.002 log2 units are therefore a null-by-construction arithmetic check, not independent evidence against compositional distortion. MOR is the informative robustness result.

The paired TRIB3-minus-DDIT4 rescue contrast was also stable: −0.524 log2CPM under total CPM and −0.525 under MOR, with bootstrap intervals excluding zero and exact one-sided *p* = 0.0143 under both normalizations. However, the disease-induction component is asymmetric. TRIB3 was strongly elevated in both independent P19 cohorts under all normalizations. DDIT4’s PERK-arm disease effect was only +0.225 log2CPM under total CPM and +0.178 under MOR, smaller than the 0.340 log2CPM median absolute cross-arm discrepancy among the 30 frozen genes; in the independent 2BAct cohort its disease effect was +0.190 under total CPM but −0.138 under MOR. DDIT4 is therefore robustly PERK-responsive but not robustly disease-elevated.

These data support two different statements. TRIB3 is both disease-elevated and PERK-responsive. DDIT4 is reproducibly suppressed by oligodendrocyte-lineage PERK deletion, but its disease induction is weak and cohort/normalization dependent. The transcriptomic rescue does not establish whether either candidate is a harmful mediator, an adaptive response, or a correlated downstream output.

### 3.4. The pharmacologic arm does not independently validate a transcript mediator

The pharmacologic arm demonstrates why the interaction term cannot stand in for normalization. Under total CPM, DDIT4 increased in treated Jimpy animals by +0.290 log2CPM while its genotype-by-treatment interaction was negative (*I* = −0.653), because DDIT4 increased even more strongly in treated WT animals. The same pattern persisted after non-myelin CPM and median-of-ratios normalization. TRIB3 showed a small negative P19 2BAct point estimate (*R*^*JP*^ *≈ −*0.11) under every normalization, but its bootstrap interval crossed zero and the exact one-sided test remained *p* = 0.457.

The paired TRIB3-minus-DDIT4 P19 2BAct contrast was −0.401 log2 units under all three normalization schemes, with bootstrap intervals spanning zero and exact one-sided *p* = 0.114. Therefore the pharmacologic arm does not resolve a transcript-specific TRIB3-versus-DDIT4 rescue effect at P19.

### 3.5. Independent-cohort calibration leaves TRIB3 ordinary and DDIT4 uninterpretable

The primary specificity calibration used the GSM-disjoint P19 PERK cohort for disease matching and the P19 2BAct cohort for rescue evaluation. TRIB3 remained essentially median under every normalization: 51.5th percentile under total CPM and the denominator control and 52.5th under non-myelin MOR. Thus, its small negative P19 2BAct response is not exceptional among genes independently matched for disease effect, abundance, and residual variability (Fig. 5; Table 3).

**Table 3.**
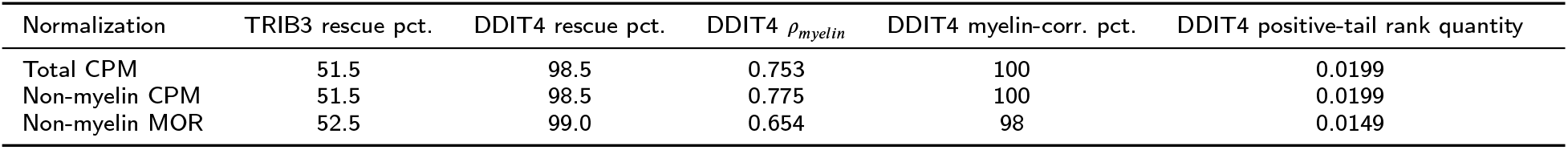
Independent-cohort P19 2BAct specificity calibration. The rescue percentile is the fraction of 200 matched genes with *R*^*JP*^ less than or equal to the candidate value. Rank quantities are descriptive, not formal *p* values.

**Figure 5.**
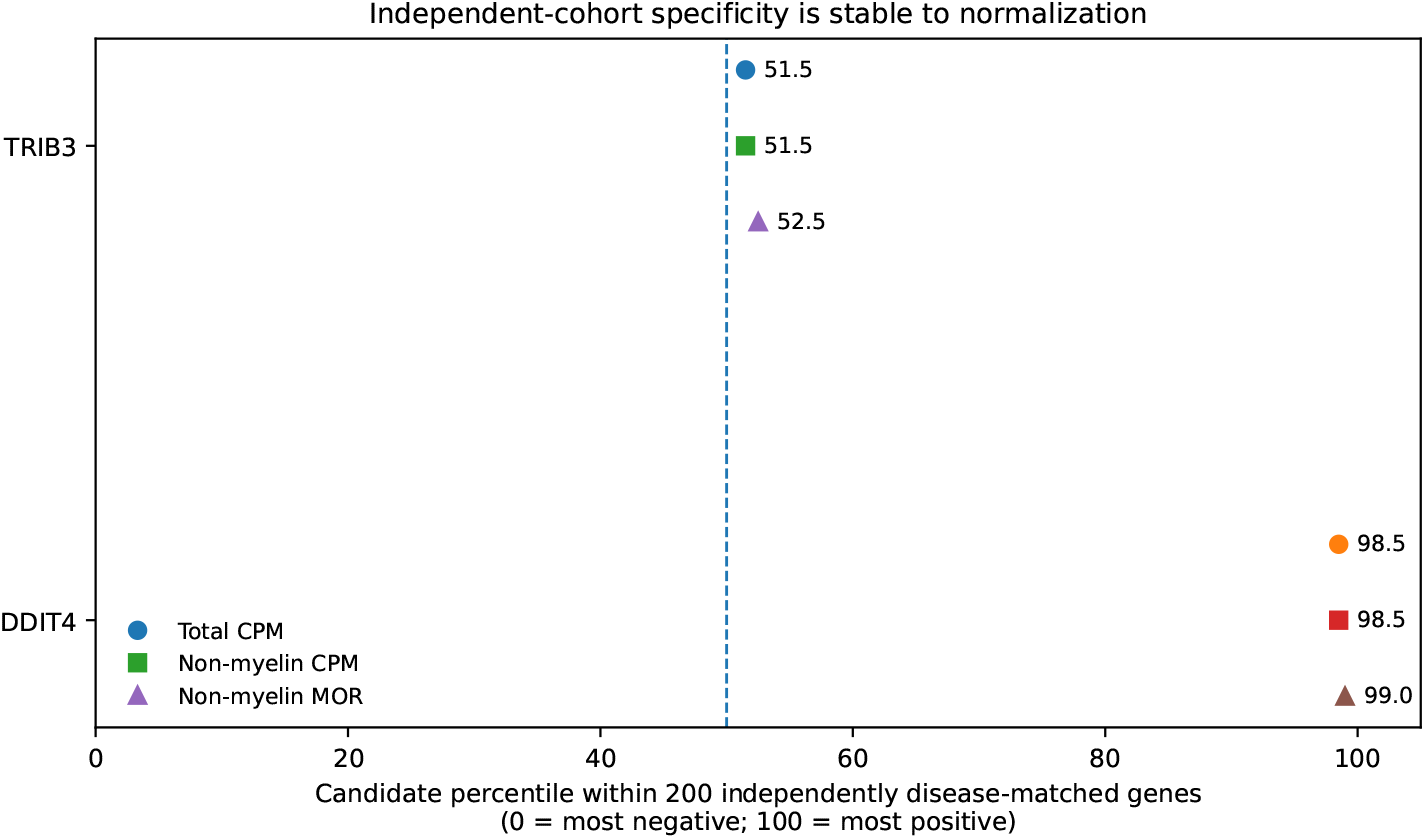
Independent-cohort specificity calibration for P19 2BAct across normalization schemes. Disease matching was defined in the disjoint P19 PERK cohort. TRIB3 remains near the center of its matched background under all normalizations. DDIT4 remains near the positive tail, but its evaluation-arm disease effect is lower than that of its independently matched background, biasing its rank upward through the same shared-control geometry. Cross-fitting is not shown because leave-one-out splitting of four untreated Jimpy controls was non-discriminating.

The direction of residual coupling bias can be assessed by comparing the disease effect in the evaluation arm with of the independently selected background. Under total CPM, TRIB3 had *D* = +1 633 in the 2BAct cohort whereas median of its 200 matched genes was +0.700; under MOR the corresponding values were +1.620 and +0.523. TRIB3 therefore carries substantially more evaluation-arm disease induction than its background and is exposed to more downward shared-control coupling. Its near-median rescue percentile is consequently conservative with respect to the conclusion that TRIB3 is not unusually normalized.

DDIT4 shows the opposite mismatch. Under total CPM its 2BAct disease effect was +0 190 while the matched-kground median was +0 430; under MOR DDIT4 was −0 138 while the background median was +0 125. DDIT4 herefore exposed to less downward coupling than the background against which its rescue percentile is ranked, biasing the percentile toward the positive tail. The 98.5–99.0th percentile cannot therefore be interpreted as evidence for an opposing biological response. The candidate-local mechanical slope implies that this mismatch contributes in the expected positive direction but does not by itself account for the full observed *R*^*JP*^.

DDIT4 also remained unusually correlated with the myelin/oligodendrocyte proxy (*ρ* = 0.75, 0.77, and 0.65), at the 100th, 100th, and 98th percentiles of its matched background. However, across matched genes the association between rescue effect and myelin-proxy correlation was weak (*ρ ≈* 0.13–0.14), so composition does not provide a demonstrated general explanation either. The positive-tail result is therefore best treated as statistically and compositionally confounded rather than mechanistically interpretable.

Leave-one-out cross-fitting was non-discriminating at four untreated-Jimpy controls. TRIB3’s four split percentiles were 2.0, 25.0, 74.0, and 96.5, with two on each side of the matched-set median. DDIT4’s were 8.0, 77.5, 90.5, and 100.0. We therefore use neither the split medians nor their superficial agreement with the independent-cohort analysis as corroborating evidence; split-level results are retained only in Supplementary Fig. S6.

### 3.6. Separate P19 cohorts constrain disease-association claims

Under total CPM, the median absolute difference in untreated Jimpy-versus-WT disease effects across the 30 frozen genes was 0.340 log2CPM, with an interquartile range of 0.216–0.571 and a 90th percentile of 0.682. Even the untreated-WT frozen myelin-set fraction differed from 0.7971% in the PERK cohort to 0.9243% in the 2BAct cohort (16.0% relative), providing a second, independent sign of between-cohort variation. The 0.340 log2CPM quantity is therefore used as an empirical working variability scale, not a formal reproducibility floor. TRIB3 differed by 0.662 log2CPM between the PERK and 2BAct cohorts (+0.971 versus +1.633) but remained strongly positive in both. DDIT4 differed by only 0.035 under total CPM (+0.225 versus +0.190), yet its 2BAct disease effect becomes −0.138 under MOR. Because DDIT4’s PERK-arm disease induction is itself only +0.178–+0.225 log2CPM, the data do not support calling DDIT4 robustly disease-associated. The original cross-arm plot is retained as Supplementary Fig. S7 strictly as a cohort-variability audit.

### 3.7. Age, composition, and transcript-program structure further limit pharmacologic interpretation

The 2BAct age series is not monotonic for TRIB3. Its Jimpy treatment effect is positive at P7 (+1.323 SD), negative at P14 (−1.052 SD), and only slightly negative at P19 (−0.143 SD). DDIT4 is positive at all three ages (+0.985, +0.449, and +0.469 SD). The P7 TRIB3 sign flip means that “TRIB3 normalizes under 2BAct” is not a defensible age-general statement; the response is state-dependent (Supplementary Fig. S2).

Whole-brain composition adds another limitation. The myelin/oligodendrocyte proxy increased under both rescues. Under P19 2BAct, the unadjusted DDIT4 point estimate changed sign in a diagnostic model containing that proxy, and TRIB3’s point estimate also moved substantially (Supplementary Fig. S3). Because the proxy is treatment-responsive, this analysis is not a causal correction. It demonstrates only that the bulk-tissue candidate contrast is sensitive to changing oligodendrocyte/myelin representation. The matched-background audit independently showed that DDIT4 itself is in the extreme positive tail of myelin-proxy correlation (Supplementary Fig. S8), although matched-gene rescue effects correlate only weakly with myelin-proxy correlation.

Previously defined stress programs also failed to behave as uniformly coherent latent variables. The p53-effector set had median pairwise Spearman correlations of −0.021 and 0.060 in the PERK and 2BAct arms, respectively; the ATF4–DDIT4 and CHOP–TRIB3 programs also changed coherence by arm. In contrast, the myelin/differentiation program was highly coherent (median *ρ* = 0.957 and 0.931; PC1 94.6% and 97.0%). Program means are therefore retained only as descriptive summaries (Supplementary Fig. S4).

The same caution applies to p53. At gene level, *Cdkn1a* decreased after PERK deletion (*R*^*JP*^ = −0.266 log2CPM, bootstrap interval [−0.407, −0.111], exact one-sided *p* = 0.0286), while *Mdm2* increased slightly and *Pmaip1* was unresolved. Under 2BAct, *Bbc3* showed a more negative and better-resolved rescue estimate than TRIB3 (*R*^*JP*^ = −0.339, bootstrap interval [−0.620, −0.020]), even though the exact one-sided test remained coarse (*p* = 0.114) and the p53 program as a whole was incoherent. That an exploratory p53 effector can look more “rescue-consistent” than the nominated TRIB3 candidate in this arm is a concrete illustration of the rule’s poor specificity. Other p53-related genes moved in heterogeneous directions, so no p53 mechanism claim is made from the composite score.

### 3.8. Supporting datasets show why induction alone is insufficient

The earlier disease-transition and pre-death datasets remain useful as orthogonal context. In GSE111595, *Ddit4* was strongly elevated in Jimpy OPCs before the differentiation-associated pathological interval, whereas several other stress and apoptosis-related transcripts did not move with it. In pre-death tunicamycin-treated OPCs (GSE81792), DDIT4, TRIB3, DDIT3, p53-related genes, and adaptive proteostasis genes could all be induced before overt cell death, while HSPA5 and HERPUD1 even moved in opposite directions at 8 h (Supplementary Fig. S1). These observations motivate perturbational adjudication but do not identify an executor.

The two human oligodendrocyte datasets are retained only as external context. Their gene-level patterns differ by cohort and lesion context, and with only 14 donor-context profiles in the correlation analysis we do not interpret failure to resolve a correlation as evidence of independence. The corrected human figure includes only GSE180759 and GSE118257 (Supplementary Fig. S5).

## 4. Discussion

### 4.1. Known statistical phenomena, rescue-specific geometry, and the transferable result

The statistical ingredients exposed here are established rather than newly discovered. Interaction evidence must be separated from simple effects Nieuwenhuis et al. (2011); baseline–change relationships can be distorted by mathematical coupling Oldham (1962); Tu and Gilthorpe (2007); RNA-seq normalization can be composition-sensitive Robinson and Oshlack (2010); Anders and Huber (2010); and cross-fitting is an established data-splitting strategy Chernozhukov et al. (2018). The transferable contribution is their specific geometry in rescue adjudication and the diagnostics that make that geometry visible before a candidate is promoted mechanistically.

The first rescue-specific failure is semantic. A genotype-by-rescue interaction answers whether treatment responses differ by genotype. It does not answer whether the disease group moved toward control. DDIT4 under P19 2BAct demonstrates this directly: the interaction is negative even though the transcript increases in treated Jimpy animals. Any mechanistic use of the word “reversal” should therefore report *R*^*JP*^ itself, with uncertainty.

The second failure is statistical. Disease induction and apparent rescue reuse the untreated-disease mean with opposite signs. Under independent groups, their covariance contains the exact term 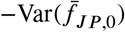. The mirror analysis shows that the post-MOR enrichment is conditional on disease-effect selection and argues against a one-sided threshold-scaling explanation, but symmetry alone cannot distinguish shared-control coupling from genuine broad disease-state restoration. The attribution to coupling comes from the slope decomposition: under MOR, the full-data observed *R*^*JP*^ -on-*D* slope is −0.256, of which the estimated mechanical term is −0.125 (48.7% of the negative slope magnitude). Deleting one of the four untreated-Jimpy samples moved the descriptive ratio from 37.8% to 90.1%; the observed-slope range was −0.341 to −0.217, and a conventional normal-theory jackknife sensitivity interval for that slope spans zero. Thus the denominator gradient being decomposed is itself not statistically resolved at this replication level. Candidate-local point estimates were 60.8% near TRIB3’s abundance and 38.6% near DDIT4’s; notably, their paired difference stayed positive in all four deletions (8.6–27.7 points), while its magnitude varied substantially. The relevant lesson is therefore not a precise mechanical fraction, but that the contrast geometry creates a potentially substantial shared-control channel whose quantitative size requires more untreated-disease replication, alongside a residual that can contain genuine biology.

The threshold-prevalence audit is therefore not an auxiliary descriptive statistic. It is a calibration test for the proposed mechanistic rule itself, and its conclusion does not depend on assigning the whole gradient to one mechanism. If broad biological restoration causes many disease-selected genes to reverse, the same rule is still too nonspecific to identify which one mediates rescue; if shared-control coupling contributes materially, the rule is additionally mechanically enriched. A candidate should not be interpreted as exceptional merely because it satisfies a criterion that classifies a large fraction of the transcriptome. Its natural companion is the arm-wide *R*^*JP*^ /*R*^*W T*^ /*I* audit, which distinguishes transcriptome-wide normalization displacement from disease-selection-conditioned enrichment.

The normalization audit adds a separate robustness lesson. The non-myelin CPM comparison is a denominator-arithmetic control whose tiny effect was predetermined by the frozen myelin set occupying less than 1.5% of reads in every P19 cell; it cannot by itself reassure against broader compositional normalization. The informative MOR sensitivity preserved the PERK candidate rescue estimates while substantially changing genome-wide threshold prevalence and DDIT4’s 2BAct disease effect. This asymmetry is structural as well as empirical: *R*^*JP*^ contains no WT samples and is insulated from WT-specific displacement, whereas *I* inherits any change in *R*^*W T*^ . Reporting *R*^*JP*^ with uncertainty is therefore not only semantically cleaner than calling *I* “normalization”; in this design it is also the estimand protected from the WT displacement that makes interaction prevalence fragile.

### 4.2. What changes biologically in PMD

The biological consequence is not the discovery of a new PMD death executor; it is a change in the evidential status of the leading downstream candidates and in what experiment should come next. Chen et al. (2026) already established TRIB3 induction in Jimpy and reduction after oligodendrocyte-specific PERK deletion, and nominated TRIB3/AKT. The present reanalysis confirms that TRIB3 is disease-elevated and genetically PERK-responsive, but it removes the P19 2BAct transcriptome as independent transcript-specific corroboration: neither TRIB3 nor DDIT4 has a resolved Jimpy rescue effect in that pharmacologic arm, and TRIB3 is ordinary relative to an independently calibrated background. The case for TRIB3 therefore remains biologically serious, but it rests on genetic PERK dependence, disease induction, prior AKT biology, and a still-needed direct necessity experiment rather than on two independently discriminating transcriptomic rescues.

DDIT4 supports a different claim. It is robustly suppressed by oligodendrocyte-lineage PERK deletion, but its disease induction is weak and not reproducible across cohort/normalization combinations. It should therefore be treated as a PERK-responsive comparator worth direct perturbation, not as a second established disease mediator. This distinction matters mechanistically because ATF4 can induce DDIT4 and REDD1 can inhibit mTORC1 Whitney et al. (2009); Kimball and Jefferson (2012); Brugarolas et al. (2004), while REDD1 can support survival in other stress contexts Dennis et al. (2013). A direct *Ddit4* perturbation coupled to oligodendrocyte survival and phospho-S6/4E-BP1 measurements would distinguish a harmful branch from an adaptive or canonical PERK output.

The broader biological message concerns experimental format. In hypomyelinating disease, successful rescue changes exactly the tissue-defining oligodendrocyte/myelin program that dominates bulk-brain composition, while the nominated stress genes can reside at very different abundance and precision regimes. Bulk rescue transcriptomics can therefore be convincing evidence that an intervention restores a disease-associated state yet remain weak evidence for which downstream transcript mediates that restoration. For TRIB3, oligodendrocyte-lineage loss or targeted suppression with AKT phosphorylation and survival outcomes would test necessity directly; for DDIT4, the corresponding mTORC1 readouts are required. More generally, cell-resolved or targeted measurements should replace further bulk ranking when the question is mediator identity rather than state restoration.

### 4.3. Why the pharmacologic arm should be interpreted differently

Six observations converge on a conservative interpretation of 2BAct. First, the total-CPM transcriptome-wide excess is driven primarily by a large positive WT treatment response, and MOR substantially recenters that displacement while leaving the Jimpy response distribution near zero. Second, after MOR the remaining enrichment is concentrated among genes selected on disease effect and reverses direction among disease-decreased genes. That mirror establishes disease-selection dependence but is compatible with both genuine broad restoration and shared-control coupling; the slope decomposition, not the symmetry alone, gives a full-data point estimate near one-half for the shared-control term, but the sample-deletion analysis shows that this fraction is poorly resolved at four untreated-Jimpy replicates. Third, neither TRIB3 nor DDIT4 has a resolved P19 disease-group rescue effect under any normalization. Fourth, independent matching places TRIB3 near the center of its background despite TRIB3 having substantially greater evaluation-arm disease induction than that background, making the ordinary rank conservative. Fifth, DDIT4’s positive-tail percentile is biased in the opposite direction because its evaluation-arm disease effect is lower than its matched background, while its disease status is itself normalization-sensitive. Sixth, age, abundance, and bulk-tissue composition remain important: TRIB3 changes sign between P7 and later ages and lies in the lowest expression decile, whereas DDIT4 is exceptionally correlated with the myelin/oligodendrocyte proxy.

The DDIT4 98.5–99.0th percentile is therefore not an interpretable positive mechanistic result. The rank-tail quantity is small, but matched genes are correlated, two candidates were examined, and the candidate/background disease-effect mismatch violates the exchangeability intuition needed to read the percentile biologically. Its extreme myelin association adds a second candidate-specific ambiguity, while the weak matched-background correlation between rescue and myelin association argues against a simple composition-only explanation.

The 2BAct arm nevertheless remains informative at the perturbation level. It supports the source study’s conclusion that altering ISR signaling improves Jimpy phenotype Chen et al. (2026); it is much weaker for ranking individual downstream transcripts in whole-brain RNA-seq. The *Bbc3* example reinforces this point: BBC3 shows a more negative and better-resolved pharmacologic rescue estimate (*R*^*JP*^ = −0.339 log2CPM, 95% bootstrap CI [−0.620, −0.020]) than TRIB3 (−0.112, [−0.750, +0.586]), even though BBC3 was only an exploratory p53 effector. In an arm where the ranking rule is demonstrably nonspecific, such a pattern cannot by itself establish mechanistic priority.

### 4.4. Implications beyond the PLP1 case

The framework applies whenever a secondary perturbation analysis uses the same disease controls to define abnormality and apparent normalization. The present case separates phenomena that can otherwise be conflated in a single high-prevalence number: normalization can displace the interaction distribution transcriptome-wide; conditioning on disease effect can produce a sign-reversing mirror whether the underlying source is mechanical coupling or genuine broad restoration; and shared-control coupling can then be quantified from the cross-gene slope rather than inferred from symmetry alone. The safeguards are therefore sequential. Report the actual disease-group treatment effect separately from interactions; inspect arm-wide *R*^*JP*^, *R*^*W T*^, and *I* distributions under a defensible normalization; use the mirror analysis to test whether enrichment follows disease-effect selection; and estimate the mechanical component with the slope decomposition only when untreated-disease replication is adequate to stabilize that estimate. Coupling should be examined in an expression-stratified or candidate-local manner rather than assuming a transcriptome-wide percentage applies uniformly.

When matching on disease severity, there is no assumption-free fix for shared-control coupling. Independent-cohort calibration breaks literal sample reuse but trades that bias for a transportability requirement: disease-effect structure must remain sufficiently comparable between matching and evaluation cohorts. TRIB3 and DDIT4 show how violations can bias candidate percentiles in opposite directions. Cross-fitting avoids cohort transfer but trades transportability for precision and becomes non-discriminating when partitions are unstable. The practical hierarchy is therefore replication-aware: threshold prevalence, arm-wide response distributions, and the mirror remain useful transcriptome-wide diagnostics at the present replication, whereas the slope decomposition and cross-fitting depend directly on the number of untreated-disease biological replicates and are not quantitatively stable at *n* = 4. These are conditional diagnostics, not universal corrections; a reader should use the low-replication diagnostics first and reserve sample-partition or slope-decomposition corrections for designs with enough untreated-disease replication to support them.

These safeguards are especially important when sample sizes are small. Exact randomization tests become coarse, bootstrap intervals can be wide, and a single untreated-disease group mean can dominate both disease and rescue contrasts. Under such conditions, the appropriate conclusion may be that an intervention changes a broad disease-associated transcriptional state without identifying a specific mediator.

### 4.5. Limitations

The framework is demonstrated in one disease system rather than benchmarked across multiple perturbation datasets. Generalization to other factorial rescue designs, continuous treatments, repeated measures, or single-cell perturbations requires external validation. The decoupling, normalization, abundance-stratified coupling, and sample-deletion uncertainty audits were added after external review of the initial completed analysis, although each correction was frozen before rerunning the relevant computation and the superseded analyses remain available for audit. With only four untreated-Jimpy samples, the delete-one ranges are necessarily coarse. Moreover, because the procedure perturbs only the untreated-disease group while holding treated and WT samples fixed, it isolates the shared-control channel rather than bounding total uncertainty in the cross-gene slope.

The independent-cohort matching strategy assumes that disease-effect structure is sufficiently transportable across the two P19 cohorts to define a meaningful background. The cross-arm audit shows that this assumption is imperfect. TRIB3 remains clearly disease-elevated but differs substantially in magnitude across cohorts; DDIT4 has only weak PERK-arm disease induction and changes sign in the 2BAct cohort under MOR. The resulting candidate/background mismatch can itself shift rescue percentiles even when the cohorts are GSM-disjoint. Cross-fitting avoids cohort transfer in principle, but at only four untreated Jimpy controls the split percentiles span almost the entire possible range and are non-discriminating. The sample size at which cross-fitting becomes useful should be established by simulation or multi-dataset benchmarking rather than inferred here.

GSE277705 is bulk whole-brain RNA-seq. The genetic intervention is oligodendrocyte-lineage-specific, which gives the PERK contrast strong biological leverage, but changing cell composition remains possible. 2BAct is systemic and produces broader tissue effects. The frozen eight-gene denominator control has little mathematical leverage and therefore cannot rule out broader compositional effects; MOR is the informative normalization sensitivity. The composition-proxy analysis remains diagnostic rather than causal because the proxy is itself treatment-responsive.

The supporting GSE111595 and GSE81792 experiments have only three biological replicates per cell and are used for temporal context rather than definitive inference. The human MS datasets are cross-sectional and are not evidence for PMD causality. Finally, no transcriptomic pattern establishes molecular necessity. Direct TRIB3 and DDIT4 perturbations with biochemical pathway readouts remain the decisive experiments.

## 5. Conclusions

Rescue perturbations can strengthen mechanistic inference only when the estimand matches the biological claim, shared-control coupling is recognized, cross-cohort calibration is audited for transportability, and normalization sensitivity is explicit. In the PLP1/Jimpy case, oligodendrocyte-specific PERK deletion robustly suppresses TRIB3 and DDIT4, but the two transcripts support different biological claims: TRIB3 is disease-elevated and PERK-responsive, whereas DDIT4 is robustly PERK-responsive but only weakly and inconsistently disease-induced. The P19 2BAct transcriptome does not provide transcript-specific evidence that TRIB3 normalizes, and DDIT4’s positive-tail rank is not interpretable because its evaluation-arm disease effect is lower than that of its matched background. Total CPM creates a transcriptome-wide negative interaction displacement driven primarily by a large positive WT response; MOR substantially removes that displacement and is primary for arm-wide inference. Under MOR, the remaining excess reverses direction with disease-effect selection, showing that the enrichment is selection-conditioned but not, by symmetry alone, whether it is mechanical or biological. The slope decomposition gives a 48.7% full-data mechanical-to-observed ratio, but its four-sample delete-one range is 37.8–90.1%; candidate-local point estimates are 60.8% near TRIB3 and 38.6% near DDIT4, and their paired difference remains positive in all four deletions (8.6–27.7 points) while its magnitude remains unstable. The observed cross-gene slope itself is not statistically resolved by four untreated-Jimpy replicates, so the decomposition should not be used here as a precise correction. More generally, the prevalence audit, arm-wide response distributions, and mirror provide low-replication diagnostics of nonspecificity and selection dependence, whereas slope decomposition and cross-fitting require substantially more untreated-disease replication. Without that hierarchy, a phenotypically successful rescue can generate a persuasive but misleading transcript ranking without supplying transcript-specific evidence of mediation, potentially redirecting downstream perturbation experiments toward the wrong mechanistic target.

## Funding

HJ’s research work was supported in part by the National Institutes of Health IDeA grant P20GM103408. AG’s research was funded in part by a grant from the National Multiple Sclerosis Society (RG-2305-41469).

## Competing interests

The authors declare no competing interests.

## Data and code availability

All source expression datasets are publicly available through NCBI GEO under GSE111595, GSE81792, GSE277705, GSE180759, and GSE118257. A complete versioned reproducibility archive is provided with this submission as Supplementary Data S1. It contains the frozen sample manifests, identifier-mapping diagnostics, original and corrected rescue classifications, bootstrap and exact-permutation outputs, independent-cohort matched backgrounds, the v0.3 decoupling analyses, the v0.4 normalization audit, the v0.5 abundance-stratified coupling audit, the v0.6 arm-wide interaction-distribution and size-factor diagnostic, the v0.7 symmetric disease-increased/disease-decreased prevalence diagnostic, the v0.10 delete-one untreated-Jimpy coupling-uncertainty audit, threshold-prevalence and shared-control-coupling outputs, program-coherence and composition-sensitivity analyses, and executable notebooks.

## Ethics statement

The study used only publicly available, de-identified transcriptomic data and did not involve new human-participant or animal experimentation by the authors.

## CRediT authorship contribution statement

**Hasan M. Jamil:** Conceptualization, Methodology, Software, Formal analysis, Investigation, Data curation, Visualization, Writing – original draft, Writing – review and editing, Funding acquisition. **Alexander Gow:** Validation, Biological interpretation, Supervision, Writing – review and editing.

## Notes

### Competing Interest Statement

The authors have declared no competing interest.

