## Supplemental figures and tables for "Why rescue-based transcript ranking can mislead: normalization displacement and shared-control coupling in perturbation transcriptomics"

### Supplementary Figure S1: temporal and pre-death context

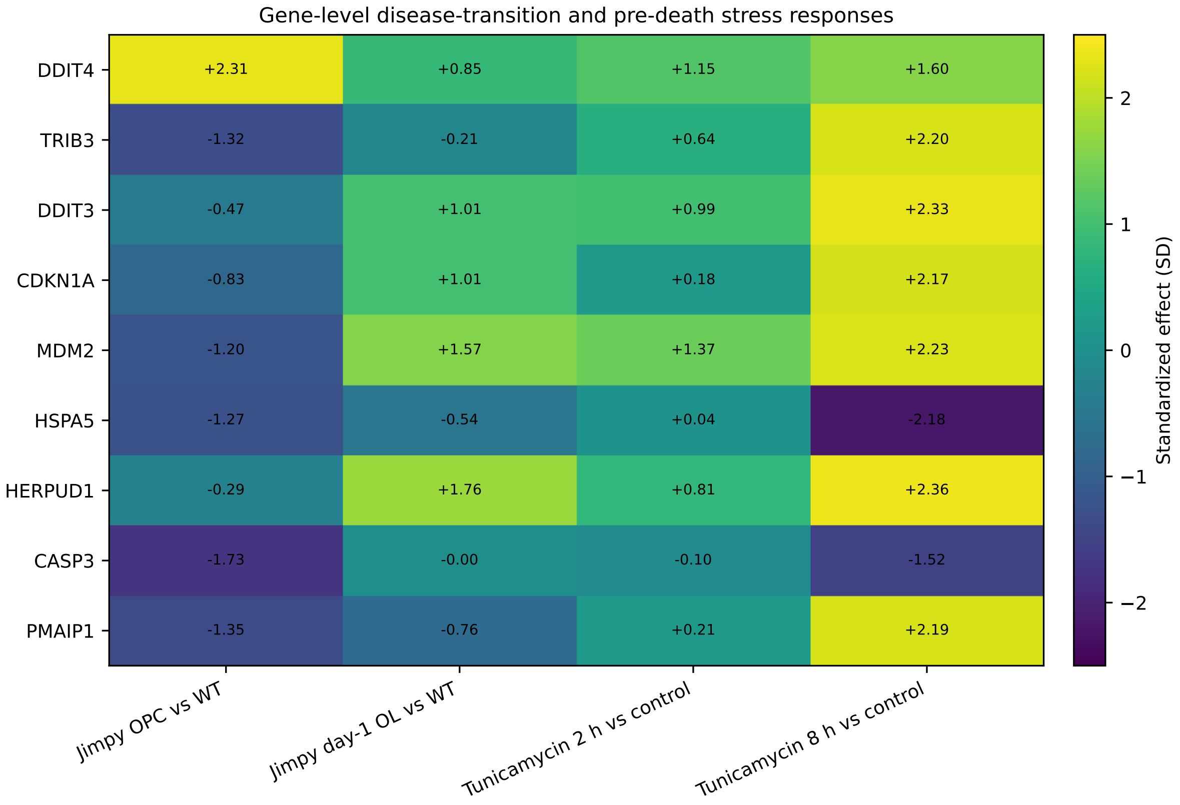


Figure 1: Gene-level standardized effects in the Jimpy differentiation experiment (GSE111595) and pre-death tunicamycin experiment (GSE81792). The supporting datasets show that stress-response genes can diverge before overt death; they are not used to identify a downstream executor.

#### Supplementary Figure S2: 2BAct age series

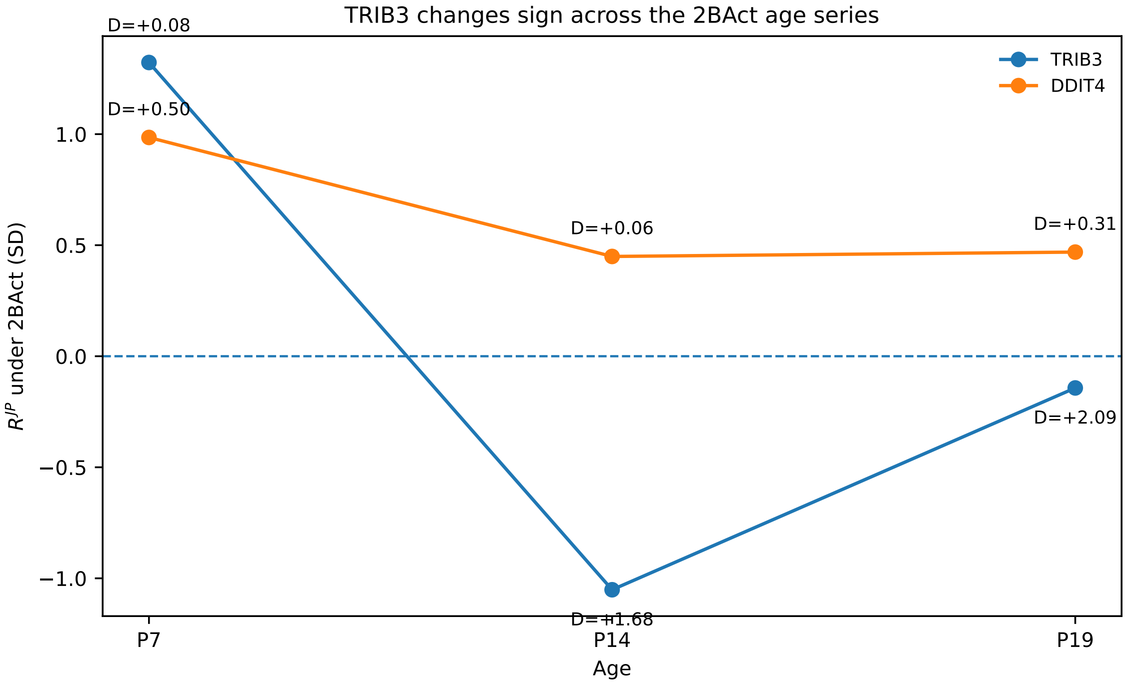


Figure 2: Descriptive age-dependent 2BAct response. TRIB3 changes from a positive Jimpy treatment effect at P7 to negative point estimates at P14 and P19. DDIT4 is positive at all three ages. Labels report the untreated Jimpy-versus-WT disease effect $D$ in SD units.

#### Supplementary Figure S3: composition sensitivity

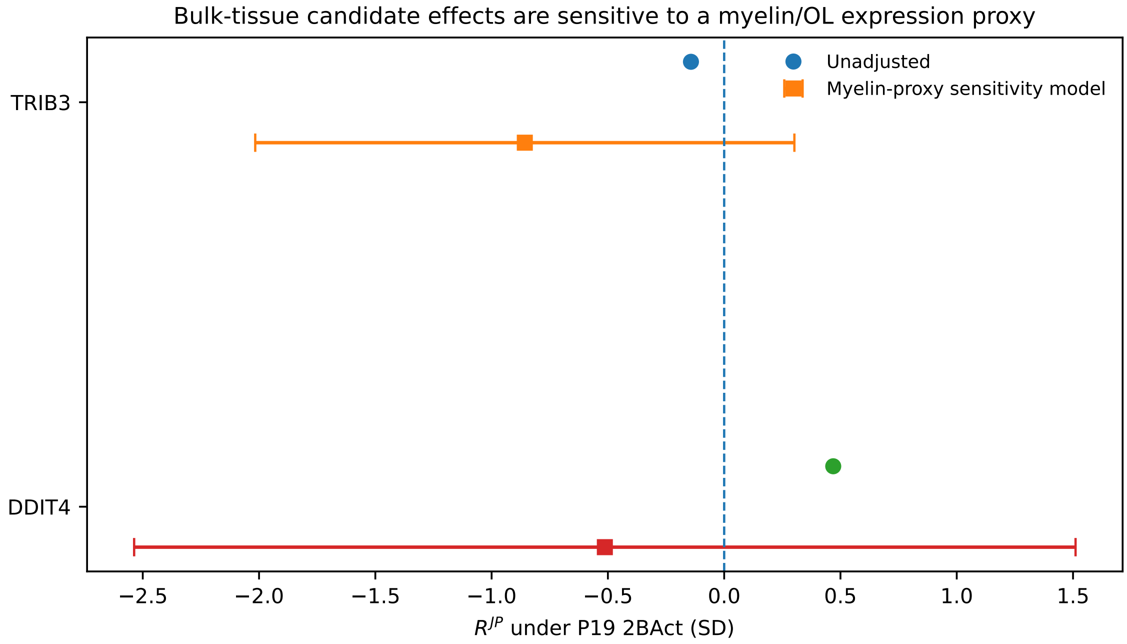


Figure 3: Diagnostic P19 2BAct composition sensitivity. The adjusted model contains a myelin/oligodendrocyte expression proxy. Because the proxy changes under rescue and may lie on the causal pathway, adjusted estimates are not interpreted causally.

#### Supplementary Figure S4: program coherence

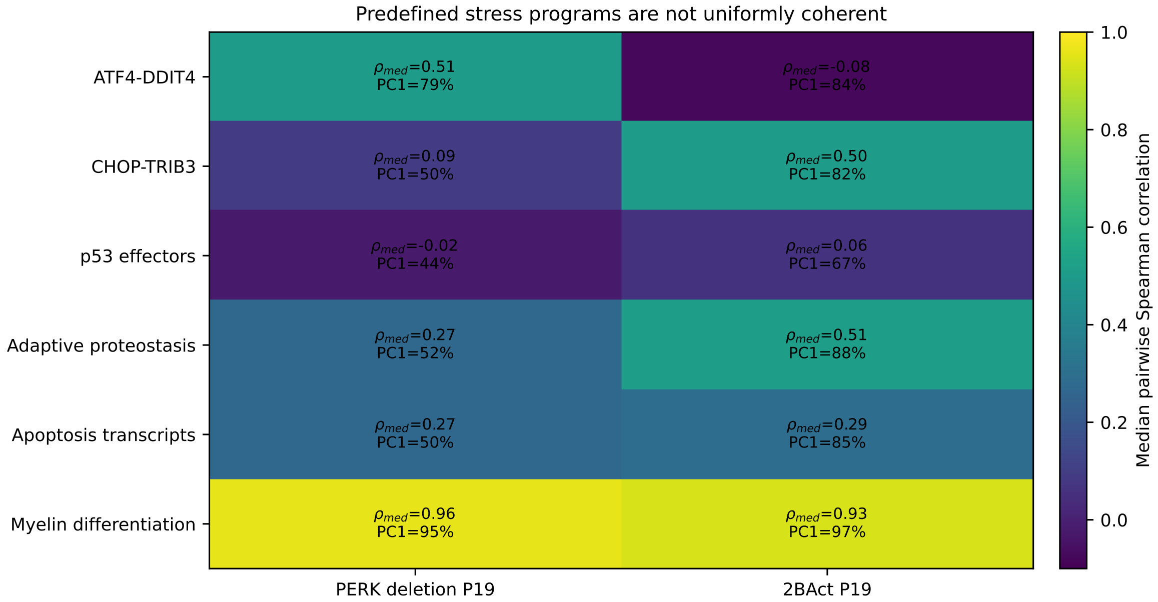


Figure 4: Internal coherence of predefined P19 programs. Myelin/differentiation genes are highly coherent in both arms, whereas several mechanistic stress programs are not. Program means are therefore descriptive rather than primary mechanistic estimands.

### Supplementary Figure S5: human oligodendrocyte context

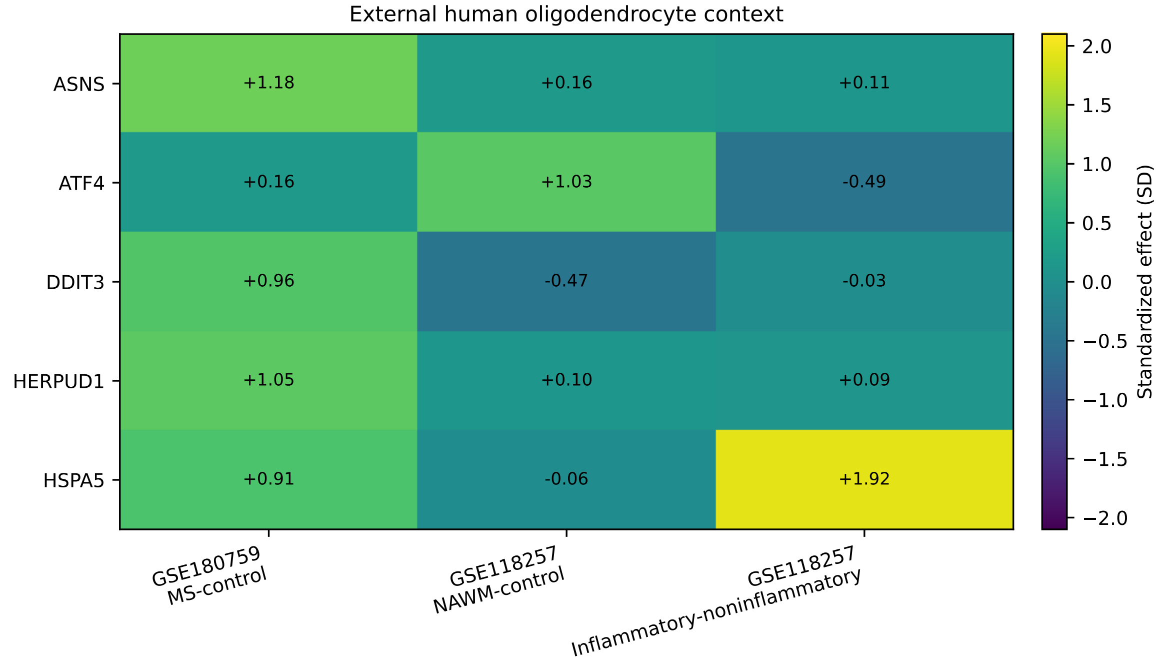


Figure 5: Corrected external human oligodendrocyte context. Only the two declared human cohorts, GSE180759 and GSE118257, are shown. These data provide context for stress-program heterogeneity and are not used as causal evidence for PMD.

#### Supplementary Figure S6: leave-one-out cross-fit

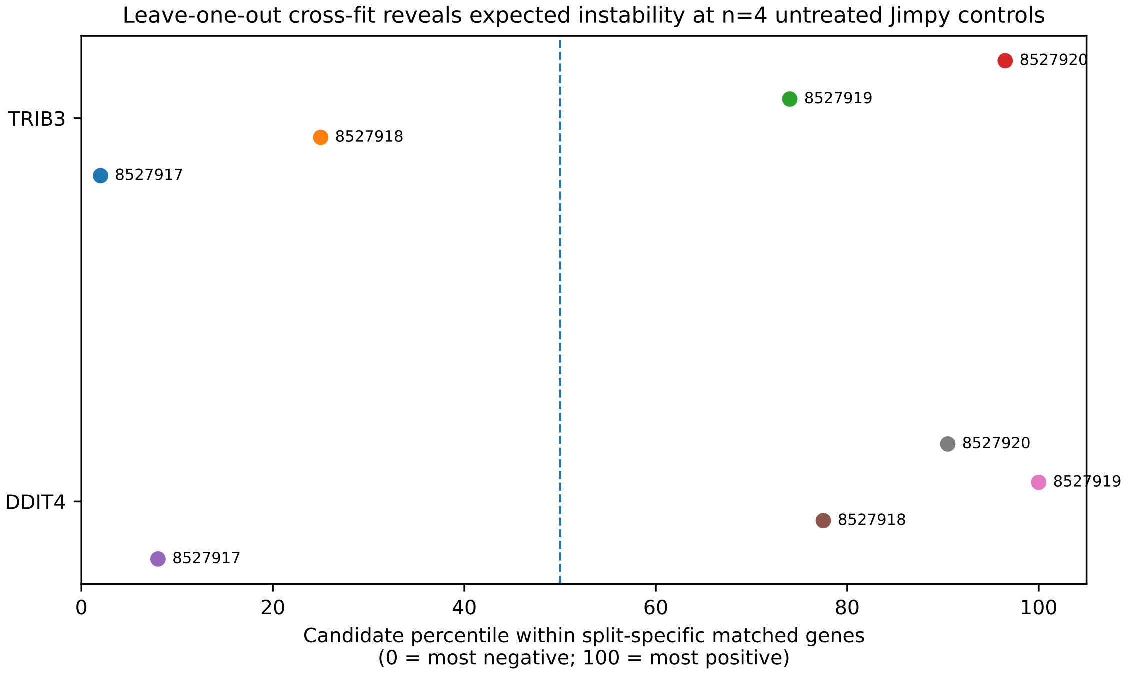


Figure 6: Split-specific candidate percentiles from leave-one-out cross-fitting of the four untreated Jimpy P19 2BAct controls. TRIB3 spans 2.0–96.5 and DDIT4 8.0–100.0. These ranges are non-discriminating; the split medians are not used as corroborating evidence.

#### Supplementary Figure S7: cross-arm disease-effect variability

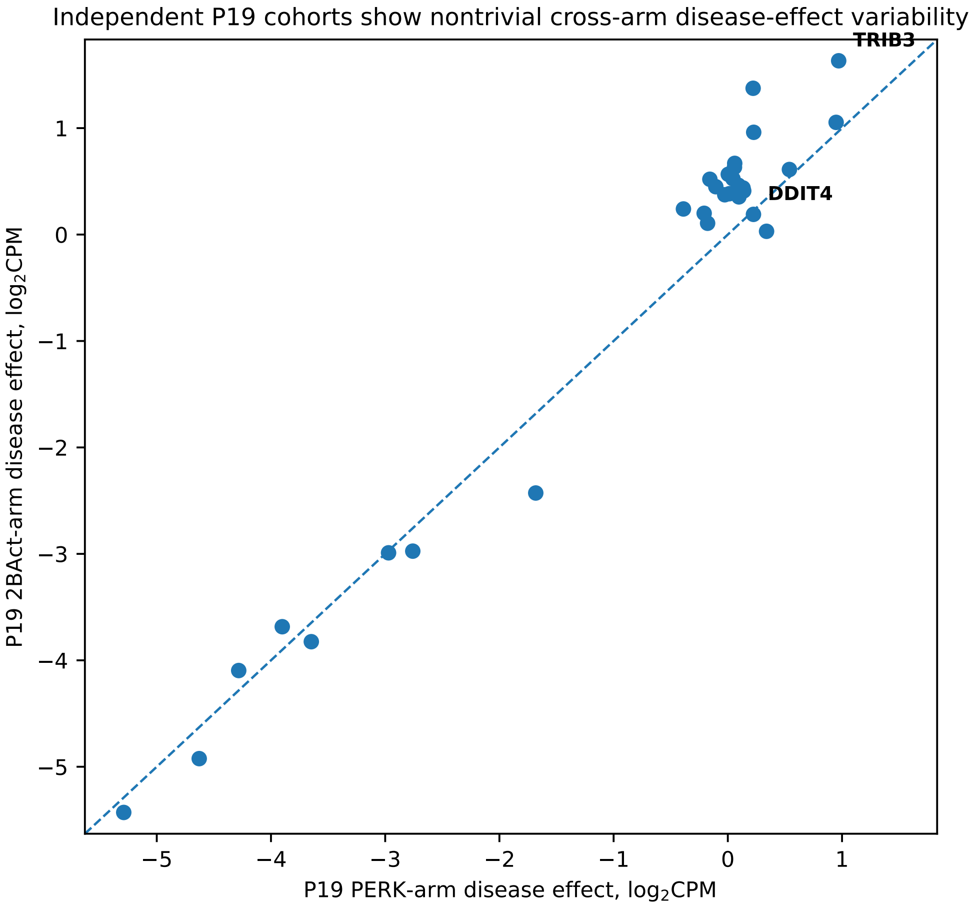


Figure 7: Total-CPM untreated Jimpy-versus-WT disease effects in the two GSM-disjoint P19 cohorts. The figure is an empirical cohort-variability audit rather than evidence of candidate reproducibility. TRIB3 remains positive in both cohorts despite a large magnitude difference; DDIT4's weak 2BAct disease effect changes sign under non-myelin median-of-ratios normalization.

#### Supplementary Figure S8: DDIT4 myelin-proxy association

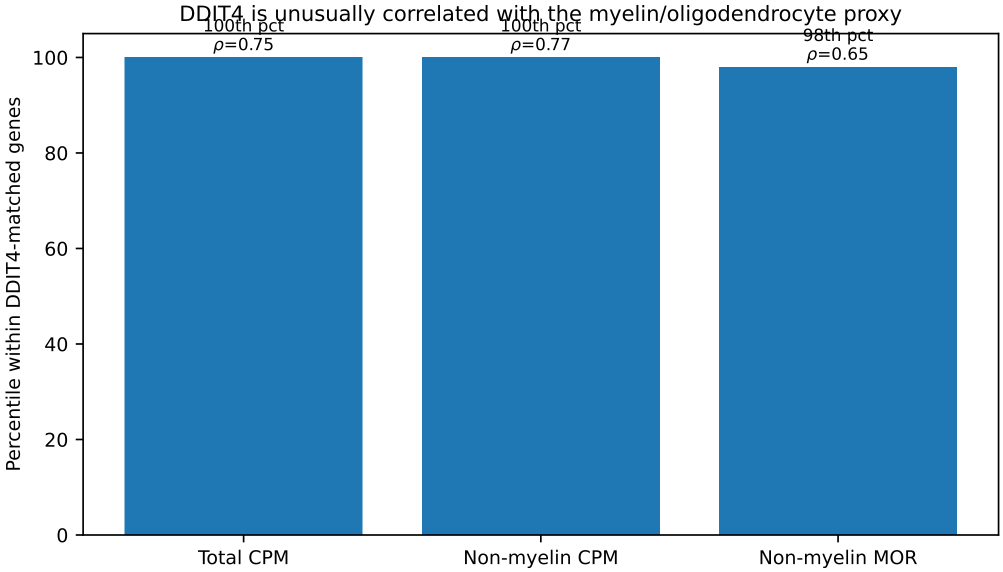


Figure 8: DDIT4's Spearman correlation with the myelin/oligodendrocyte proxy expressed as a percentile of its 200 independently disease-matched genes. DDIT4 is at the 100th, 100th, and 98th percentiles under total CPM, the denominator control, and non-myelin MOR, respectively. Across matched genes, rescue response and myelin-proxy correlation are only weakly associated, so this extreme candidate-level correlation is a caution rather than a demonstrated causal explanation.

#### Supplementary Table S1: normalization and coupling audit

Table 1: P19 2BAct normalization sensitivity and shared-control coupling.

| Normalization | All genes $I \leq -0.25$ | Disease-eligible $I \leq -0.25$ | Strict rule | Mechanical slope | Observed slope |
|---|---|---|---|---|---|
| Total CPM | 75.6% | 85.9% | 54.3% | -0.142 | -0.291 |
| Non-myelin CPM | 76.0% | 86.6% | 54.7% | -0.142 | -0.290 |
| Non-myelin MOR | 47.6% | 77.6% | 58.8% | -0.125 | -0.256 |

The frozen eight-gene myelin-set fractions of total reads were, respectively, 0.7971%, 0.9774%, 0.0838%, and 0.2040% for PERK WT control, WT rescue, Jimpy control, and Jimpy rescue; the corresponding 2BAct values were 0.9243%, 1.4633%, 0.0838%, and 0.1211%. Excluding the set from the CPM denominator changes the Jimpy rescue contrast by only 0.00174 log2 units in PERK and 0.00054 in 2BAct, so that normalization is an arithmetic leverage control rather than an informative alternative size-factor model.

#### Supplementary Table S1B: abundance-stratified coupling audit

Table 2: P19 2BAct shared-control coupling under non-myelin MOR. The ratio is the descriptive mechanical-to-observed negative-slope point estimate; sample-deletion sensitivity is reported in Supplementary Table S1F.

| Expression decile | Median baseline log-expression | Mechanical slope | Ratio |
|---|---|---|---|
| 1 | 0.558 | -0.153 | 67.8% |
| 2 | 1.162 | -0.137 | 54.2% |
| 3 | 1.858 | -0.142 | 49.1% |
| 4 | 2.654 | -0.127 | 40.8% |
| 5 | 3.444 | -0.123 | 42.4% |
| 6 | 4.169 | -0.111 | 41.7% |
| 7 | 4.919 | -0.094 | 37.1% |
| 8 | 5.646 | -0.116 | 48.0% |
| 9 | 6.497 | -0.093 | 43.8% |
| 10 | 7.759 | -0.057 | 26.8% |

Candidate-local 500-gene abundance neighborhoods gave full-data ratios of 60.8% for TRIB3 (expression decile 1) and 38.6% for DDIT4 (decile 8). Their paired difference is not resolved by the four-sample delete-one sensitivity analysis (Supplementary Table S1F). These values are descriptive finite-sample diagnostics, not causal decompositions.

#### Supplementary Table S1C: genome-wide response distributions

Table 3: Genome-wide medians and interquartile ranges of $R^{JP}$, $R^{WT}$, and $I$ for both P19 rescue arms under all normalization schemes. Effects are in within-arm gene-standardized SD units. Each column is summarized marginally across genes; therefore median $I$ need not equal median $R^{JP}$ minus median $R^{WT}$ even though $I = R^{JP} - R^{WT}$ holds gene by gene.

| Arm | Normalization | $R^{JP}$ median [IQR] | $R^{WT}$ median [IQR] | $I$ median [IQR] | $I < 0$ |
|---|---|---|---|---|---|
| PERK | Total CPM | +0.016 [−0.452, +0.474] | +0.562 [−0.006, +1.035] | −0.525 [−1.168, +0.202] | 69.1% |
| PERK | Non-myelin CPM | +0.023 [−0.442, +0.479] | +0.573 [+0.007, +1.043] | −0.527 [−1.172, +0.197] | 69.2% |
| PERK | Non-myelin MOR | +0.032 [−0.479, +0.535] | +0.036 [−0.649, +0.631] | +0.049 [−0.710, +0.796] | 48.2% |
| 2BAct | Total CPM | −0.002 [−0.261, +0.338] | +1.016 [+0.493, +1.348] | −1.014 [−1.476, −0.278] | 81.2% |
| 2BAct | Non-myelin CPM | −0.000 [−0.259, +0.339] | +1.040 [+0.517, +1.367] | −1.035 [−1.494, −0.297] | 81.6% |
| 2BAct | Non-myelin MOR | +0.003 [−0.435, +0.464] | +0.154 [−0.720, +0.940] | −0.149 [−1.141, +0.946] | 53.4% |

#### Supplementary Table S1D: library sizes and MOR size factors

Table 4: Cell-level raw-library-size and median-of-ratios (MOR) size-factor summaries. Values are medians; myelin percentages are cell means.

| Arm | Cell | $n$ | Raw reads | MOR factor | Myelin reads |
|---|---|---|---|---|---|
| PERK | JP control | 4 | 21,350,796 | 0.917 | 0.0838% |
| PERK | JP rescue | 4 | 21,795,362 | 0.948 | 0.2040% |
| PERK | WT control | 4 | 26,195,176 | 1.131 | 0.7971% |
| PERK | WT rescue | 4 | 19,696,426 | 0.936 | 0.9774% |
| 2BAct | JP control | 4 | 22,582,142 | 1.149 | 0.0838% |
| 2BAct | JP rescue | 3 | 23,744,270 | 1.182 | 0.1211% |
| 2BAct | WT control | 3 | 25,298,582 | 1.276 | 0.9243% |
| 2BAct | WT rescue | 3 | 26,791,390 | 1.368 | 1.4633% |

The two untreated-Jimpy cell means that both round to 0.0838% are not duplicates: direct raw-count recomputation gives 0.083776% for the PERK cohort and 0.083822% for the GSM-disjoint 2BAct cohort. The agreement at four decimal percentage points is coincidental.

### Supplementary Table S1E: symmetric prevalence diagnostic under P19 2BAct MOR

Table 5: Mirror-image prevalence after selecting genes on disease effect under non-myelin MOR. Disease-increased genes use the frozen negative-interaction rules; disease-decreased genes use sign-reversed mirror rules. The near-identical strict-rule prevalence shows that the enrichment reverses with disease-effect selection; this symmetry is compatible with both shared-control coupling and genuine broad disease-state restoration and is not, by itself, a mechanism-specific diagnostic.

| Criterion | $D \geq +0.25$ | $D \leq -0.25$ | Mirror rule for $D \leq -0.25$ |
|---|---|---|---|
| $n$ selected | 8,335 | 6,806 | – |
| Interaction threshold | 77.6% | 74.0% | $I \geq +0.25$ |
| Interaction + reversal fraction | 77.1% | 72.8% | $I \geq +0.25$, $-I/D \geq 0.25$ |
| Strict direction-consistent rule | 58.85% | 58.77% | above and $R^{JP} > 0$ |

The same within-arm z standardization is applied to both disease-effect selections. The reversal of interaction enrichment with the sign of $D$ therefore argues against a one-sided threshold-scaling explanation and shows that the enrichment is conditional on disease-effect selection. It does not uniquely attribute that enrichment to shared-control coupling, because genuine broad reversal of the disease state predicts the same sign pattern. Mechanistic attribution to coupling instead uses the cross-gene slope decomposition reported in Supplementary Tables S1 and S1B.

#### Supplementary Table S1F: sample-deletion sensitivity of coupling estimates

Table 6: Delete-one untreated-Jimpy sensitivity analysis for P19 2BAct under non-myelin MOR. Each of the four untreated-Jimpy biological samples was deleted in turn while the treated-Jimpy and WT groups were held fixed; MOR normalization was re-estimated and the coupling quantities were recomputed on the frozen gene sets. The leave-one-out range is the primary sensitivity summary. It isolates perturbation of the shared untreated-disease term rather than bounding total sampling uncertainty in the slope. Ratios and ratio differences are shown in percentage points.

| Estimand | Full-data estimate | Leave-one-out range |
|---|---|---|
| Transcriptome mechanical slope | −0.125 | [−0.202, −0.129] |
| Transcriptome observed slope | −0.256 | [−0.341, −0.217] |
| Transcriptome coupling ratio | 48.7% | 37.8–90.1% |
| TRIB3-local coupling ratio | 60.8% | 46.8–89.6% |
| DDIT4-local coupling ratio | 38.6% | 28.9–78.8% |
| TRIB3 minus DDIT4 local ratio | 22.2 points | 8.6–27.7 points |

Because genes within a sample share normalization factors, a naive bootstrap over genes would understate dependence. The primary sensitivity analysis therefore deletes an entire untreated-Jimpy biological sample and recomputes MOR normalization. All four paired TRIB3-minus-DDIT4 differences remain positive, but their magnitude varies from 8.6 to 27.7 points, so the ordering is sign-stable without being quantitatively resolved. For the unbounded observed slope, a conventional normal-theory jackknife sensitivity interval is $[-0.589, +0.077]$; with only four untreated-Jimpy samples we use this only to flag that the population-level denominator gradient is not statistically resolved, not as the primary uncertainty bound. Because the treated groups are held fixed, the delete-one analysis specifically probes the shared-control channel and does not represent total slope uncertainty.

#### Supplementary Table S2: individual p53-related genes

Table 7: Gene-level p53-related rescue effects on the log2CPM scale. The heterogeneous directions do not support treating the six genes as a coherent mechanistic module.

| Arm | Gene | $D$ | $R^{JP}$ | $I$ | 95% CI for $R^{JP}$ | Exact $p_{\downarrow}$ |
|---|---|---|---|---|---|---|
| PERK deletion | CDKN1A | 0.539 | -0.266 | -0.084 | [-0.407,-0.111] | 0.0286 |
| PERK deletion | MDM2 | -0.177 | +0.067 | -0.030 | [-0.002,+0.142] | 0.9143 |
| PERK deletion | PMAIP1 | -0.105 | -0.125 | +0.372 | [-0.392,+0.114] | 0.2714 |
| PERK deletion | GADD45A | +0.339 | +0.079 | -0.067 | [-0.147,+0.343] | 0.7143 |
| PERK deletion | BBC3 | +0.222 | +0.037 | -0.144 | [-0.273,+0.359] | 0.5857 |
| PERK deletion | BAX | +0.058 | +0.235 | -0.034 | [+0.011,+0.444] | 0.9429 |
| 2BAct | CDKN1A | +0.611 | -0.183 | -0.736 | [-0.378,+0.013] | 0.0857 |
| 2BAct | MDM2 | +0.106 | +0.019 | -0.167 | [-0.069,+0.118] | 0.6000 |
| 2BAct | PMAIP1 | +0.449 | +0.228 | +0.114 | [-0.139,+0.564] | 0.8000 |
| 2BAct | GADD45A | +0.030 | +0.149 | -0.127 | [-0.077,+0.351] | 0.8857 |
| 2BAct | BBC3 | +1.374 | -0.339 | -1.242 | [-0.620,-0.020] | 0.1143 |
| 2BAct | BAX | +0.632 | -0.051 | -0.774 | [-0.185,+0.070] | 0.3714 |

#### Supplementary Table S3: P19 sample-size audit

| Experiment | Genotype | Condition | Age | $n$ |
|---|---|---|---|---|
| PERK | WT | Cre− | P19 | 4 |
| PERK | WT | Cre+ | P19 | 4 |
| PERK | Jimpy | Cre− | P19 | 4 |
| PERK | Jimpy | Cre+ | P19 | 4 |
| 2BAct | WT | control diet | P19 | 3 |
| 2BAct | WT | 2BAct | P19 | 3 |
| 2BAct | Jimpy | control diet | P19 | 4 |
| 2BAct | Jimpy | 2BAct | P19 | 3 |

Thirty individual frozen genes entered the gene-level Benjamini–Hochberg interaction correction in each P19 arm. The 16 P19 PERK samples (GSM8527867–GSM8527882) and 13 P19 2BAct samples (GSM8527911–GSM8527923) share zero GSM accessions.

#### Supplementary note: superseded within-arm matched-$D$ analysis

An earlier robustness analysis selected genes for disease-effect similarity and evaluated rescue using the same P19 2BAct untreated-Jimpy group. That analysis was retained for audit but removed from inference after recognizing that $D = \bar{f}_{JP,0} - \bar{f}_{WT,0}$ and $R^{JP} = \bar{f}_{JP,1} - \bar{f}_{JP,0}$ share $\bar{f}_{JP,0}$ with opposite signs. The final manuscript therefore uses independent-cohort disease matching as primary. Leave-one-out cross-fitting at four untreated Jimpy controls is reported only as a failed stress test of sample splitting and is not used as corroborating evidence.
